# Mechanisms that ensure speed and fidelity in eukaryotic translation termination

**DOI:** 10.1101/2021.04.01.438116

**Authors:** Michael R. Lawson, Laura N. Lessen, Jinfan Wang, Arjun Prabhakar, Nicholas C. Corsepius, Rachel Green, Joseph D. Puglisi

## Abstract

Translation termination, which liberates a nascent polypeptide from the ribosome specifically at stop codons, must occur accurately and rapidly. We established single-molecule fluorescence assays to track the dynamics of ribosomes and two requisite release factors (eRF1 and eRF3) throughout termination using an *in vitro*-reconstituted yeast translation system. We found that the two eukaryotic release factors bind together to recognize stop codons rapidly and elicit termination via a tightly regulated, multi-step process that resembles tRNA selection during translation elongation. Because the release factors are conserved from yeast to humans, the molecular events that underlie yeast translation termination are likely broadly fundamental to eukaryotic protein synthesis.

**One Sentence Summary:** Direct visualization of eukaryotic translation reveals the dynamics of termination at stop codons.

## Main Text

Protein synthesis concludes when a translating ribosome encounters a stop codon at the end of an open reading frame, triggering recruitment of two factors to liberate the nascent polypeptide: <u>e</u>ukaryotic <u>R</u>elease <u>F</u>actor <u>1</u> (eRF1), a tRNA-shaped protein that decodes the stop codon and cleaves the peptidyl-tRNA bond (*1-3*), and <u>e</u>ukaryotic <u>R</u>elease <u>F</u>actor <u>3</u> (eRF3), a GTPase that promotes eRF1 action (*4-6*). After translation termination, the ribosome, P-site tRNA, and mRNA are released via recycling (*4, 7, 8*). Despite decades of study, the order and timing of molecular events that drive translation termination remain unclear, as multistep processes are difficult to assess using traditional approaches. A cohesive understanding of translation termination and its underlying steps, central to normal translation, would also support the treatment of diseases caused by premature stop codons, which include cystic fibrosis, muscular dystrophy, and hereditary cancers (*9*). Indeed, since premature stop codons cause 11% of all heritable human diseases (*10*), stop-codon readthrough therapeutics have immense clinical potential (*9, 11*).

## Results

Here we used an *in vitro*-reconstituted yeast translation system (*12*) and single-molecule fluorescence spectroscopy to track eukaryotic release factor dynamics and termination directly. We reasoned that ribosomes translating mRNAs with very short open reading frames would provide the simplest system for detailed analysis of the discrete sub-steps of termination. Ribosome complexes were programmed with Met (M-Stop) or Met-Phe (M-F-Stop) mRNAs and reacted with saturating amounts of eRF1 and eRF3. Peptide release from both M-Stop and M-F-Stop ribosome complexes occurred at similar rates as a longer tetrapeptide (M-F-K-K-Stop) programmed ribosome complex (**Fig. 1A** and **S1A-B**), and also matched the rate previously characterized for tripeptide programmed ribosome complexes (*4, 5*). To monitor eRF1 and eRF3 binding to ribosomes in real time, we labeled both proteins specifically with fluorescent dyes (**Fig. S1C-D**) and established that both labeled proteins exhibited wild-type peptide release activity (**Fig. 1B** and **S1E-F**). Association of eRF1 with the ribosome was monitored by <u>F</u>örster <u>R</u>esonance <u>E</u>nergy <u>T</u>ransfer (FRET) between 60S subunits labeled with Cy3 (FRET donor) on the C-terminus of uL18 and eRF1 labeled with Cy5 (FRET acceptor) on the N-terminus (*13*). Structural models place these termini ∼50 Å apart when eRF1 is bound in the A site (**Fig. 1C**) (*3, 13, 14*). Next, Cy3-labeled ribosomal complexes programmed with Met in the P site and either UAA or UUC in the A site were combined with Cy5-eRF1 and unlabeled eRF3, and FRET was monitored at equilibrium using <u>T</u>otal Internal <u>R</u>eflection <u>F</u>luorescence (TIRF) Microscopy. eRF1-60S FRET was only observed when a stop codon was in the A site (**Fig. 1D** and **S1G-H**), demonstrating the specificity of the FRET signal for proper eRF1 association.

**Fig. 1:**
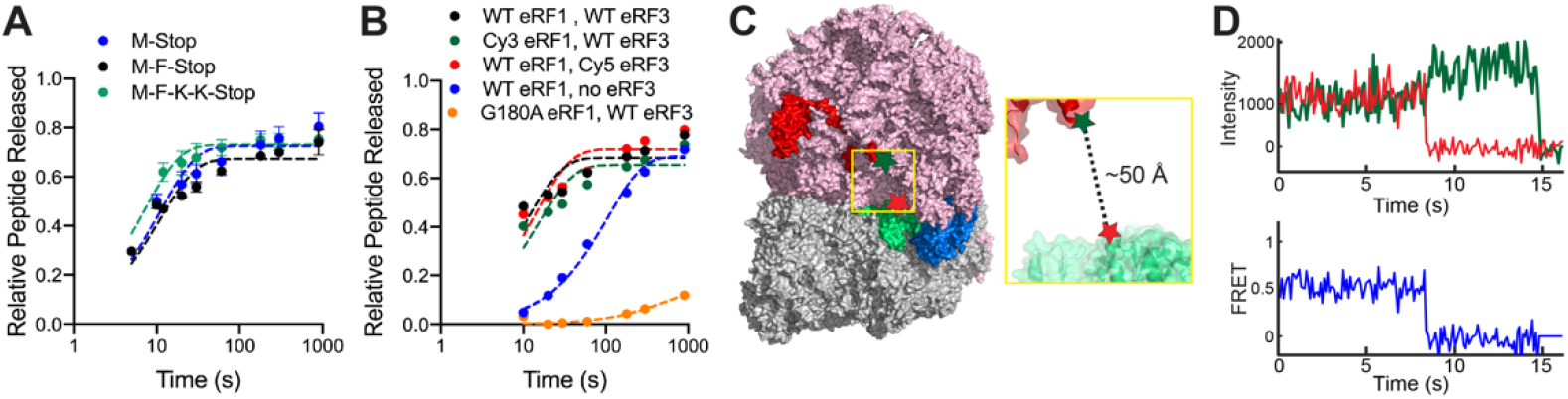
Single-molecule studies of termination. (**A**) Peptides are liberated at similar rates from ribosomes translating a variety of model mRNAs. (**B**) Wild-type and labeled release factors liberate peptides from ribosomes; catalytically-dead eRF1 (orange) is inactive. (**C**) Structural modeling (PDB ID: 5LZT (*14*)) suggests labeled eRF1 (green, Cy5-labeled at red star) would FRET with ribosomes (red, Cy3-labeled at green star) upon binding to A site. (**D**) FRET observed with Cy3-eRF1 and Cy5-60S by TIRF microscopy.

We leveraged this FRET-based binding signal to probe the roles of eRF1 and eRF3 in translation termination. We first prepared 80S ribosomal complexes programmed on 5’-biotinylated M-Stop mRNAs with Cy5-labeled 60S subunits; these complexes were tethered to neutravidin-coated <u>Z</u>ero-<u>M</u>ode <u>W</u>aveguide (ZMW) surfaces (*15, 16*). Upon start of real-time data acquisition, Cy3-eRF1, excess GTP, and unlabeled eRF3 were added to the surface, and Cy3-eRF1 and Cy5-60S fluorescence within individual ZMWs were monitored by excitation with a 532-nm laser (**Fig. 2A**).

**Fig. 2:**
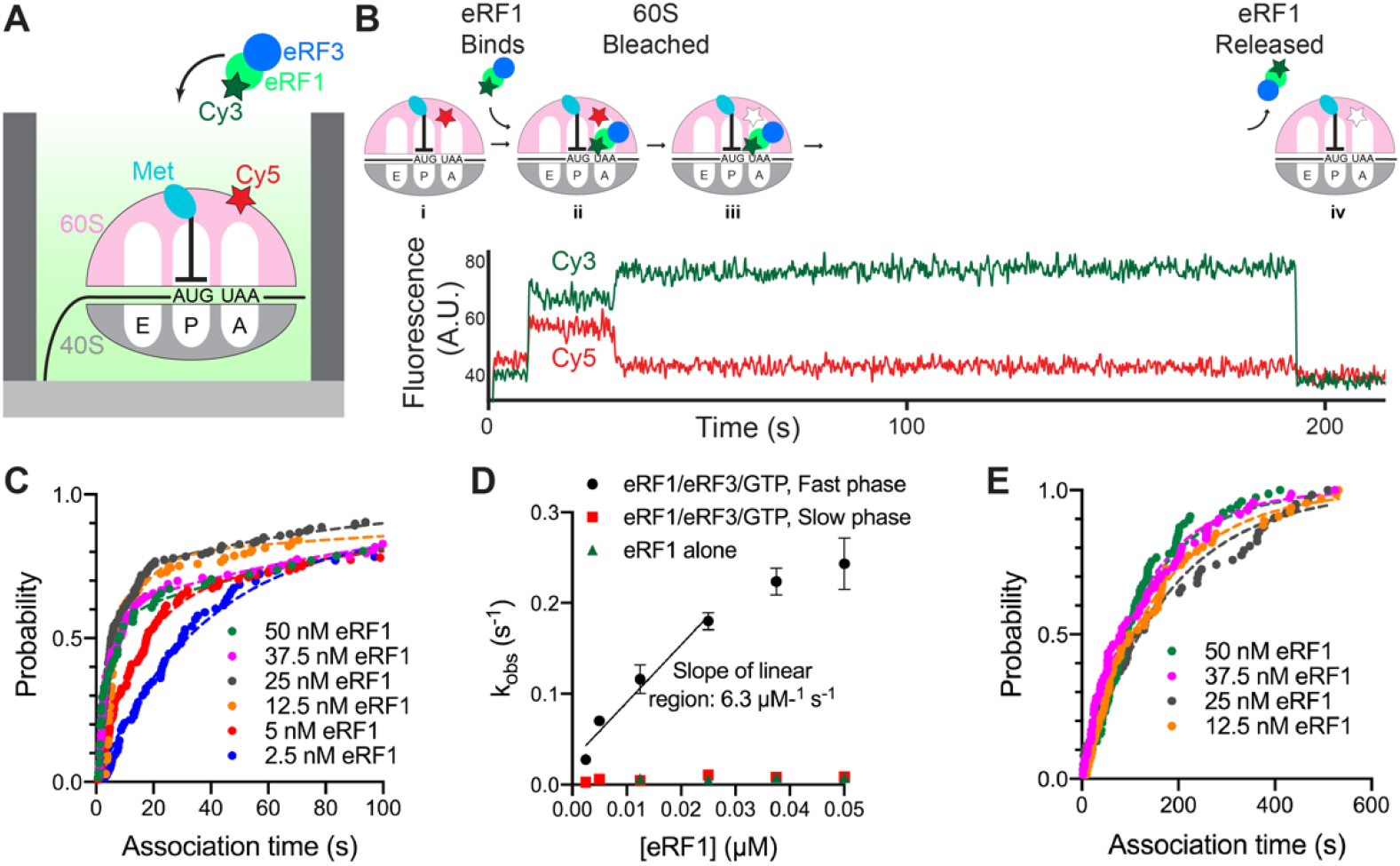
eRF3 promotes fast binding of eRF1 to ribosomes halted at stop codons. (**A**) Assay schematic. (**B**) Example of fast binding of eRF1 (Cy3, green) to M-Stop ribosomes (Cy5, red) observed in the presence of eRF3. (**C**) Binding of eRF1 to M-Stop ribosomes is fast and concentration-dependent in the presence of eRF3; arrival time distributions were fit to a double-exponential model. (**D**) Observed rates of eRF1 binding to M-Stop ribosomes (k_obs_) with and without eRF3. (**E**) Binding of eRF1 to M-Stop ribosomes is slow and eRF1 concentration-independent without eRF3; arrival time distributions were fit to an exponential model.

Rapid, concentration-dependent eRF1 binding to the ribosomal A site was detected upon delivery of the release factors (**Fig. 2B** and **S2A-C**). Association kinetics were fit to a double-exponential function with a dominant (56 – 83%) eRF1 concentration-dependent fast phase with a pseudo-first order rate constant of 6.3 ± 3.9 µM^−1^ s^−1^ (**Fig. 2C-D** and **S2B**); a minor (17 – 44%) slow phase, which did not vary with eRF1 concentration, was also observed (e.g. k_obs_ = ∼0.009 s^−1^; **Fig. 2D** and **S2B**). Conversely, eRF1 bound very slowly to these same complexes in the absence of eRF3 (e.g. k_obs_ = ∼0.008 s^−1^; **Fig. 2D-E** and **S2D-E**); the rate constant for this eRF3-independent binding was similar to the slow phase observed with eRF3, and also was unaffected by eRF1 concentration. In all cases, eRF1 binding events were long-lived (e.g. τ = 227 ± 13 s, **Fig. S2B** and **S2E**), and prolonged detection is likely limited by dye lifetime (**Fig. S2F**). Notably, in the presence of eRF3, the rapid eRF1 binding observed here is similar to the rate of Phe-tRNA^Phe^ ternary complex binding to its cognate A-site codon under similar conditions (9.0 ± 0.4 µM^−1^ s^−1^; **Fig. S2G**). These results indicate that eRF1 binding, which would otherwise be limited by a slow event, is rapid enough to compete with tRNAs for A-site occupancy when assisted by eRF3.

We next tracked eRF3 dynamics directly, independent of eRF1, to establish a baseline understanding of its interaction with the ribosome. We used a previously established inter-ribosomal subunit FRET signal to confirm 80S complex formation (*13*), and monitored dye-labeled eRF3 dynamics via fluorescent bursts that occur upon factor binding to immobilized ribosomes. Ribosomes, Cy3-labeled on uL18 and Cy5-labeled on uS19 (yielding FRET upon 80S formation), were programmed with 5’-biotinylated M-Stop mRNAs and tethered to ZMWs. Next, Cy5-eRF3 and GTP were added to ZMWs and were illuminated with 532- and 642-nm lasers. After an initial phase of FRET, typified by rapid 40S-Cy5 photobleaching, brief bursts of additional Cy5 signal were observed that marked binding and dissociation of eRF3 (**Fig. 3A**). eRF3 binding was concentration-dependent (**Fig. 3B**), and association kinetics were fit to an exponential function with a pseudo-first order rate constant of 0.4 ± 0.2 µM^−1^ s^−1^ (**Fig. S3A-B**). eRF3 resided briefly on the ribosome (τ = 0.15 ± 0.01 s; **Fig. 3C**), and the dwell times between eRF3 binding events varied with its concentration (**Fig. S3B**), consistent with a bimolecular association reaction. Inclusion of either GTP analogs or a GTPase-deficient eRF3 mutant (H348E (*4*)) did not impact its association or dissociation rates (**Fig. 3C** and **S3C**) suggesting that this binding cycle occurs independent of GTP hydrolysis.

**Fig. 3:**
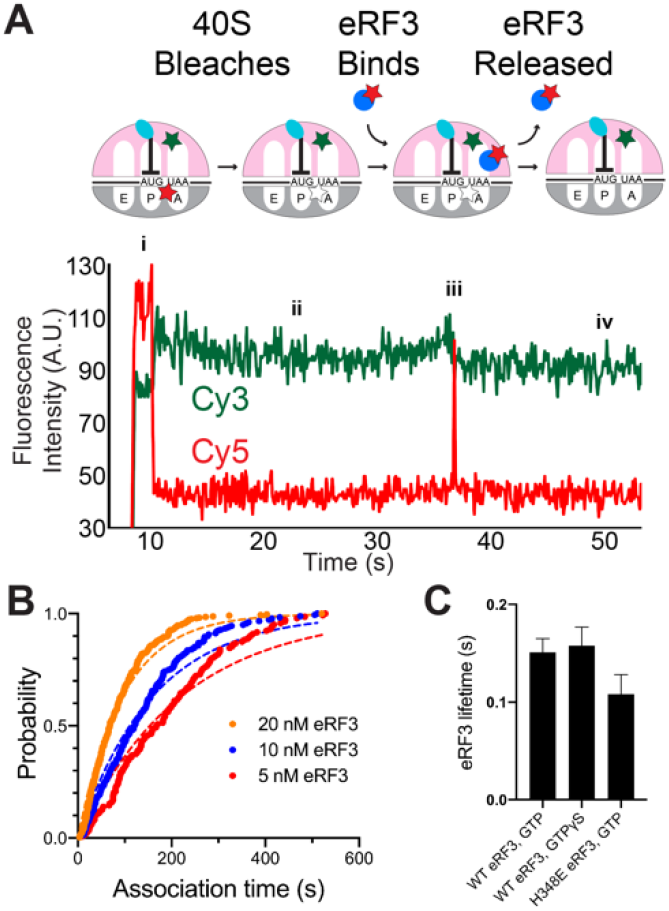
Observing eRF3 dynamics in ZMWs. (**A**) Example of eRF3 binding to M-Stop ribosomes. (**B**) Binding of eRF3 to M-Stop ribosomes is concentration-dependent; arrival time distributions were fit to an exponential model. (**C**) GTP hydrolysis by eRF3 is not required for its release from the ribosome in the absence of eRF1.

Two distinct models could explain how eRF3 promotes fast association of eRF1 with ribosomes halted at stop codons. eRF3 may first bind to ribosomes, triggering rearrangements that favor subsequent association of eRF1 with ribosomes as suggested recently (*17*). Alternatively, eRF3 may act as a chaperone, directly delivering eRF1 to ribosomes (*5*). To distinguish between these models, we performed single-molecule experiments similar to those described above but now simultaneously tracking fluorescent eRF1 and eRF3. We observed concurrent binding of the two factors to M-Stop ribosomes (**Fig. 4A**). While we also observed eRF1 and/or eRF3 binding individually to ribosomes in these experiments–unsurprising since the release factors can each bind alone to ribosomes and are at sub-saturating concentrations–the frequency of such independent binding events was very low (<0.1%), allowing us to rule out that co-arrivals occur primarily by random chance (see Methods). Analysis of Cy5 (eRF1) and Cy3.5 (eRF3) fluorescence intensities, aligned to the beginning of apparent co-association events (“post-synchronization”), further demonstrates that eRF3 binding to a stop codon-halted ribosome is transient, whereas eRF1 resides longer on the ribosome (**Fig. 4B**). Omission of GTP decreased the number of observed co-binding events by 17-fold, confirming that eRF1, eRF3, and GTP bind the ribosome together as a pre-formed ternary complex (**Fig. S4A**). Ternary complex association kinetics were fit to a double-exponential function, yielding a dominant fast phase rate that was dependent upon eRF1 concentration (**Fig. 4C** and **S4B-D**). In contrast to the dynamics of eRF3 in absence of eRF1 (where eRF3 lifetime had little dependence on GTP hydrolysis, ranging from 0.11 – 0.16 s; **Fig 3C**), here GTP accelerated eRF3 release from ribosomes by 8-fold compared to experiments performed with the more slowly hydrolyzed analog GTPγS (0.3 ± 0.1 s with GTP vs. 2.5 ± 0.1 s with GTPγS; **Fig. 4D** and **S4E-F**). Substitution of wild-type eRF3 with a GTPase-deficient mutant similarly slowed its release from the ribosome by 5-fold (1.6 ± 0.1 s; **Fig. 4D** and **S4E-F**). Thus, eRF3 is a chaperone that delivers eRF1 to ribosomes halted at stop codons, and eRF3 departure from the ribosome is partly governed by its GTPase activity.

**Fig. 4:**
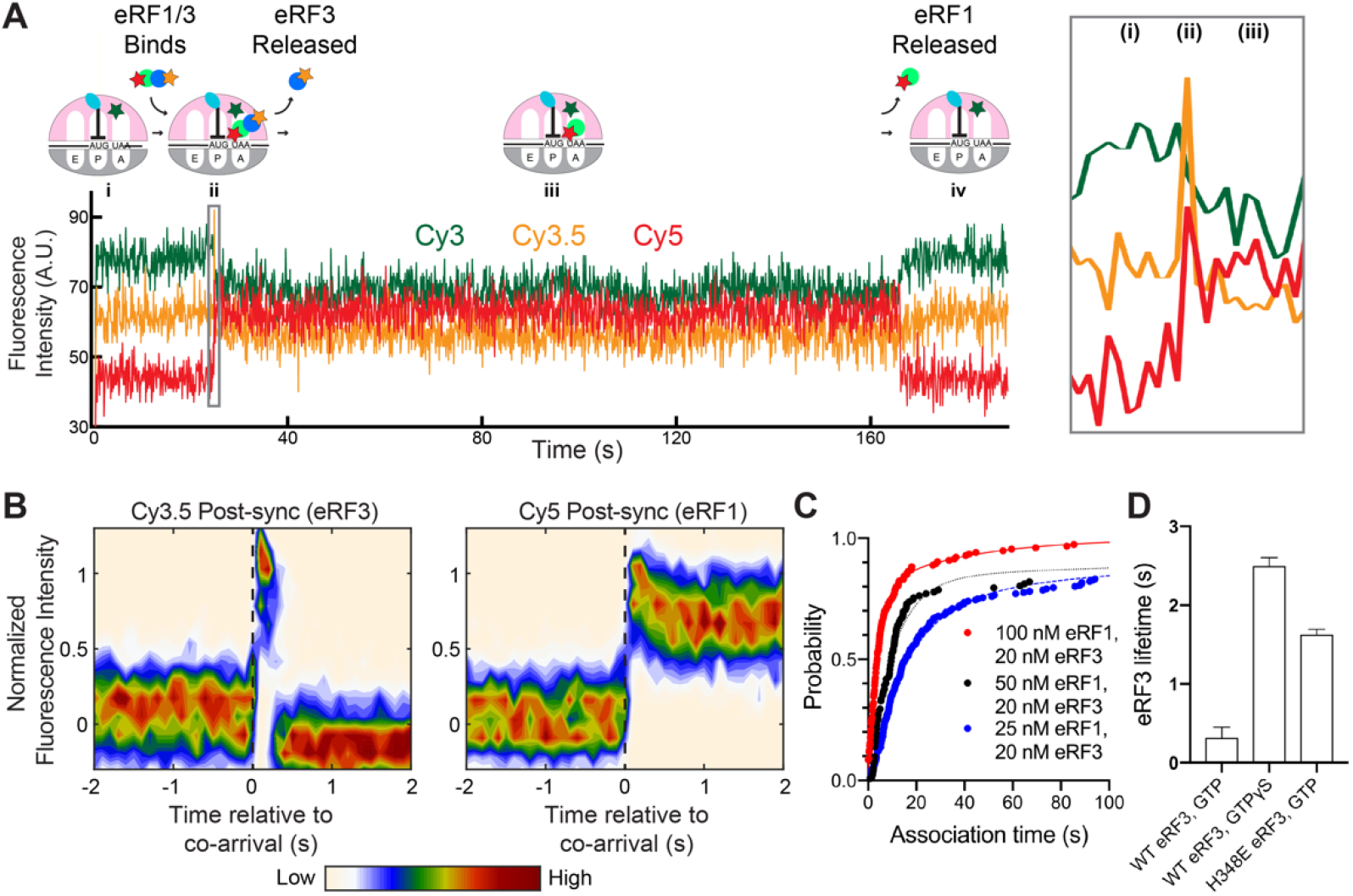
eRF3 delivers eRF1 quickly to ribosomes halted at stop codons. (**A**) Example of simultaneous binding of eRF1 (Cy5, red) and eRF3 (Cy3.5, yellow) to M-Stop ribosomes (Cy3, green). (**B**) Post-synchronization plot of fluorescence changes observed upon simultaneous binding of eRF1 and eRF3 (denoted as a dashed, black vertical line). (**C**) Simultaneous binding of eRF1 and eRF3 to M-Stop ribosomes is fast and concentration-dependent; arrival time distributions were fit to a double-exponential model. (**D**) GTP hydrolysis by eRF3 accelerates its release from the ribosome in the presence of eRF1.

We next sought to understand the timing and regulation of peptidyl-tRNA ester bond hydrolysis catalyzed by eRF1. Peptide hydrolysis triggers rapid rearrangement of P-site tRNA from a classical (P/P) to a hybrid (P/E) state (*18*), and we hypothesized this rearrangement could be tracked via FRET between labeled P-site tRNA and A-site-bound eRF1 (∼50 Å separation before vs. ∼70 Å after rearrangement; **Fig. 5A**). To test this, we tethered ribosomes programmed on an M-Stop mRNA with Cy3-labeled Met-tRNA^i^ (FRET donor) to ZMWs, added catalytically-inactive Cy5-eRF1 (G180A, FRET acceptor), unlabeled eRF3, and GTP, and illuminated with a 532-nm laser. As expected, high FRET was observed between the classical-state tRNA and eRF1 (μ = 0.632 ± 0.001; **Fig. 5B**). Next, we repeated the assay but added puromycin, a drug that cleaves the peptidyl-tRNA bond, and indeed observed lower efficiency FRET between the now-hybrid-state tRNA and catalytically-inactive eRF1 (μ = 0.531 ± 0.005; **Fig. 5B**). Therefore, peptidyl-tRNA bond status can be deduced by monitoring P-site tRNA conformation through this FRET signal.

**Fig. 5:**
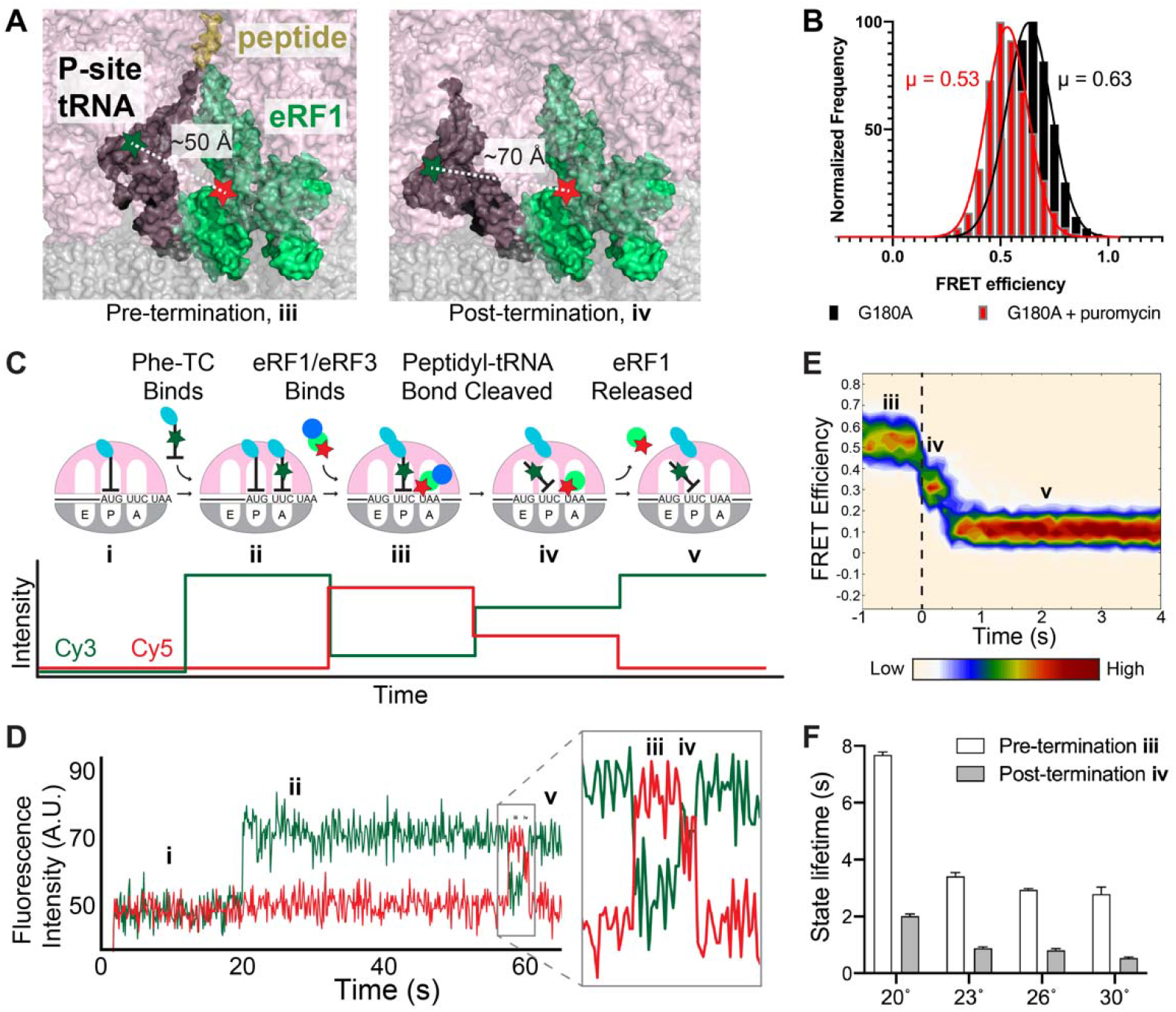
Tracking peptidyl-tRNA bond hydrolysis. (**A**) Model of FRET between eRF1 (Cy5, red star) and P-site tRNA (Cy3, green star) before and after termination. *Left*, pre-termination (modeled by 40S alignment, PDB IDs: 5LZU and 5LZT (*14*)). *Right*, post-termination (modeled by 40S alignment, PDB IDs: 5LZU and 3J77 (*14, 37*)). (**B**) FRET observed between P-site Cy3-Met-tRNA^i^ and G180A Cy5-eRF1 (black); addition of puromycin (red) yields lower FRET as expected. (**C**) Assay schematic and expected results. (**D**) Example of FRET observed with Cy3-Phe (green) and Cy5-eRF1 (red). (**E**) Post-synchronization plot of FRET efficiency observed before and after peptidyl-tRNA bond hydrolysis (marked by a dashed, black vertical line). (**F**) Pre- and post-termination state lifetimes observed at different temperatures.

We used this FRET signal to correlate eRF1 dynamics with peptidyl-tRNA bond hydrolysis in real time. Ribosomes programmed on 5’-biotinylated M-F-Stop mRNAs were tethered to ZMWs and illuminated with a 532-nm laser. We then added a mixture of Cy3-labeled Phe-tRNA^Phe^ (FRET donor), Cy5-eRF1 (FRET acceptor), excess eRF3, elongation factors, and eIF5A (an accessory factor that accelerates both elongation and termination (*19*)). Fast tRNA binding, denoted by high Cy3 signal, was observed soon after factor addition and persisted until translocation and subsequent eRF1 binding occurred (**Fig. 5C-D** and **S5A**). Association of eRF1 initially resulted in a high-FRET signal (μ = 0.6736 ± 0.003, herein referenced as “pre-termination state”; **Fig. S5B-C**) and was followed by a lower-FRET signal (μ = 0.4613 ± 0.007, “post-termination state”; **Fig. 5C-E** and **S5B**). Finally, eRF1 was released from the ribosome, resulting in restoration of high Cy3 signal (**Fig. 5C-E**). Critically, substitution of eRF1 with an inactive mutant reduced the number of observed high-to lower-FRET transitions by ∼7-fold (**Fig. S5D-E**), which closely matched the relative extent of peptide release observed in bulk with either wild-type or G180A eRF1 (**Fig. 1B** and **S1E-F**). FRET is also specific for stop codon recognition, as replacement of the UAA stop codon with near-cognate UAU completely eliminated these FRET transitions (**Fig. S5D**).

We used this eRF1/tRNA FRET assay to characterize the kinetics of peptidyl-tRNA bond hydrolysis, focusing on ribosomal events before and after termination. Pre-termination state lifetimes of eRF1 on the 80S ribosome fit well to a two-step, irreversible kinetic model with termination occurring in 2.8 s at 30 °C (95% CI: 2.6 – 3.0 s; **Fig. 5F** and **S5F-G**). eRF1 dissociation kinetics, fit to a single step model by a single exponential, revealed that eRF1 is released quickly after termination in 0.5 s (95% CI: 0.5 – 0.6 s, **Fig. 5F** and **S5G-H**); eRF1 lifetime was unaffected by laser power variation, showing that its lifetime is not limited by dye photobleaching **Fig. S5I**). Further support that peptidyl-tRNA bond hydrolysis favors eRF1 release was observed with catalytically-inactive eRF1, which resided 6-fold longer on ribosomes (**Fig. S5J**). Peptidyl-tRNA bond cleavage also hindered rebinding of eRF1 (**Fig. S5K**), demonstrating that termination decreases the affinity of eRF1 for ribosomes. The long eRF1 lifetime observed in our prior measurements (**Fig. 2** and **4**) is attributable to differences in detection methods and the omission of eIF5A from those assays, as eIF5A does not affect eRF1 association rates but does enhance termination and eRF1 release rates (**Fig. S6**) (*19*). Peptidyl-tRNA bond hydrolysis rates increased with temperature from 20 – 30 °C (**Fig. 5F, S6G**, and **S7A-C**), and subsequent Eyring and Gibbs analyses revealed that termination is regulated by a step with a significant energetic barrier (**Fig. S7D**); consistent with this notion, prior structural work demonstrated that eRF1 undergoes a large-scale conformational change (termed “accommodation”) to render it catalytically active (*3, 14*). Release of eRF1 from the ribosomal A site is also energetically costly (**Fig. 5F** and **S7E**), likely due to extensive interactions that anchor eRF1 to the stop codon (*3*). Termination proceeded at a similar rate when the UAA stop codon was changed to either UAG or UGA (**Fig. S8**), suggesting that a common mechanism is employed at all three stop codons. We therefore observed an ordered series of events at stop codons, with eRF1 eliciting peptidyl-tRNA bond cleavage within ∼3 s of ribosomal association, followed by rapid eRF1 release.

We next explored the impact of cis-acting mRNA elements on termination. Prior work uncovered a class of 3’ <u>U</u>ntranslated region (“UTR”) mRNA sequences that promote stop-codon readthrough (*20, 21*), yet it was unclear if these elements function in part by inhibiting termination. Indeed, insertion of a 6-nucleotide sequence that promotes stop-codon readthrough with 30% efficiency (CAAUUA) into the 3’ UTR of M-F-Stop mRNAs lengthened pre-termination state duration by 2-fold (5.9 s, 95% CI: 5.2 –7.0 s with CAAUUA vs. 2.9 s, 95% CI: 2.7 – 3.0 s without CAAUUA; **Fig. 6A** and **S9A-B**). Insertion of other sequences that promote readthrough at lower efficiencies also lengthened the pre-termination state, albeit to lesser degrees (**Fig. 6A** and **S9A-B**). Thus, sequences that enhance stop-codon readthrough hinder peptidyl-tRNA bond cleavage by eRF1.

**Fig. 6:**
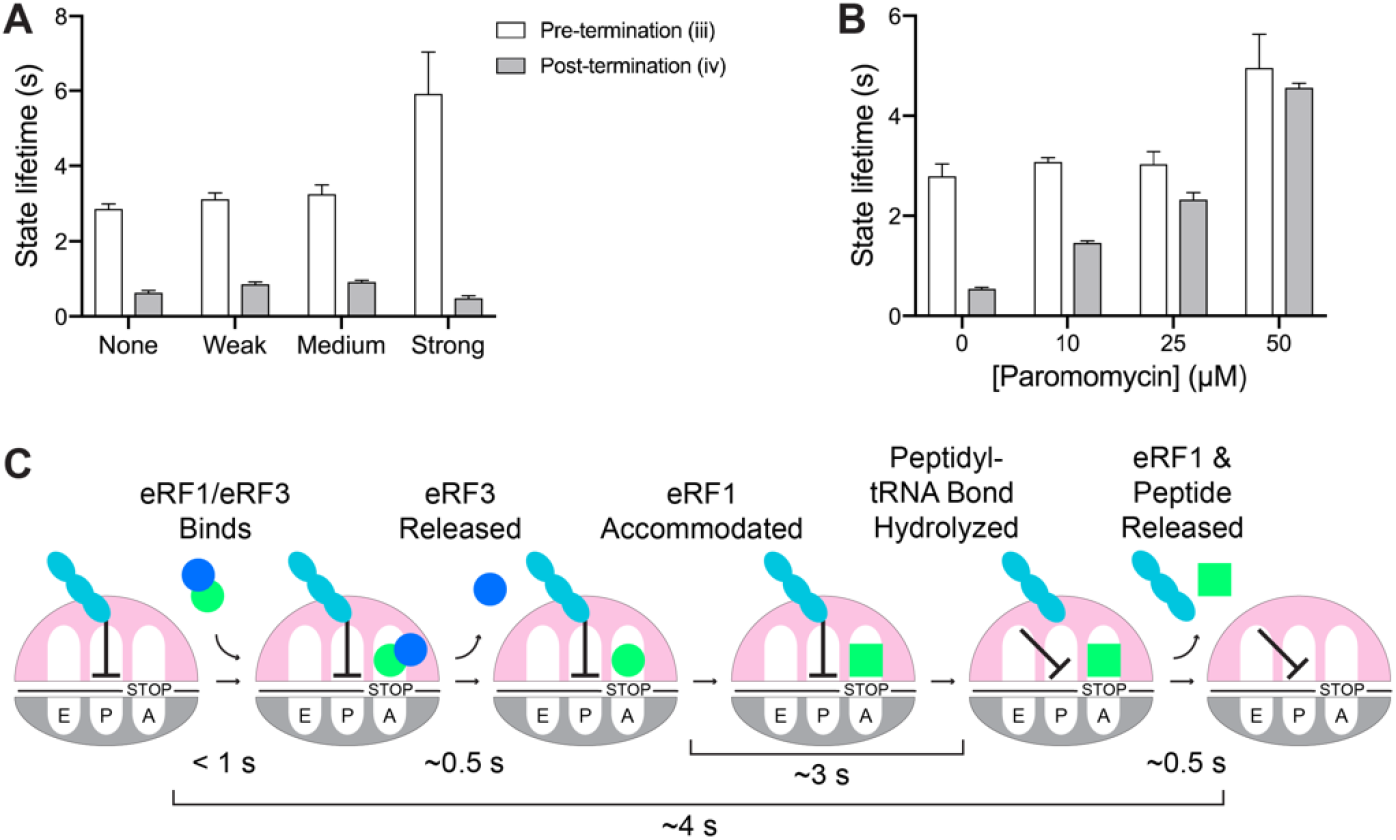
Termination is regulated by release factors, 3’ UTR mRNA sequence, and small molecules. (**A**) Insertion of 3’ UTR mRNA sequences known to promote stop-codon readthrough (“Weak” = CAAAGA, 10% efficiency; “Medium” = CAAUCA, 20% efficiency; “Strong” = CAAUUA, 30% efficiency) hinder termination; eRF1 release is unaffected. (**B**) The aminoglycoside paromomycin slows both termination and eRF1 release. (**C**) Order and timing of events in eukaryotic termination.

The aminoglycosides G418 and paromomycin also promote stop-codon readthrough (*22*), primarily by stabilizing near-cognate tRNA in the A site. Paromomycin also inhibits bacterial termination (*23*), yet its effects on eukaryotic termination were not deeply characterized (*24*). We therefore applied our suite of assays to examine the effects of these drugs on eukaryotic termination. While the addition of 50 µM G418 to our single-molecule termination assays had no discernable impact on termination kinetics (**Fig. S9C-E**), 50 µM paromomycin lengthened the pre-termination state duration (by nearly 2-fold; **Fig. 6B** and **S9C-E**), and increased post-termination state duration (by 5-fold; **Fig. 6B** and **S9C-E**). A titration of paromomycin revealed concentration-dependent effects of the drug on post-termination state duration (**Fig. 6B** and **S9C-E**). Additionally, simultaneous tracking of eRF1/3 dynamics (as described in **Fig. 4**) with 50 µM paromomycin revealed that the drug also slows eRF1/eRF3 co-binding to the ribosome (by more than 2-fold; **Fig. S9F**). Together, these studies demonstrate that stop-codon readthrough effectors hinder numerous facets of termination, thus uncovering additional nodes to target with potential therapeutics.

## Discussion

Whereas prior work broadly described the roles of eRF1 and eRF3 in regulating eukaryotic termination (*2, 4, 5, 19, 25*), here we directly monitored the kinetics of individual sub-steps to obtain a higher-resolution view of this essential process (**Fig. 6C**). First, a pre-bound ternary complex of eRF1, eRF3, and GTP rapidly binds to a stop codon-halted ribosome (**Fig. 2** and **4**). eRF3 appears to unlock eRF1 conformation to facilitate this fast ribosomal binding, as the association of eRF1 alone is slow and governed by an eRF1 concentration-independent event (**Fig. 2D-E**); consistent with this notion, prior structural studies demonstrated that the predominant conformation of eRF1 free in solution is incompatible with ribosomal binding (**Fig. S10**) (*3, 14, 26, 27*). Next, eRF3 hydrolyzes GTP to promote its own release (**Fig. 4D**), which permits the rearrangement of eRF1 to an active conformation (*14*). Accommodated eRF1 then rapidly cleaves the peptidyl-tRNA bond, triggering the ejection of both eRF1 and the liberated peptide (**Fig. 5**). Direct tracking of peptidyl-tRNA bond hydrolysis further uncovered how small molecules and mRNA sequences inhibit discrete steps in termination to promote stop-codon readthrough (**Fig. 6A-B**). Since the release factors are widely conserved from yeast to humans (**Fig. S11**) (*1, 6*), we propose that the termination mechanisms described here are fundamental to eukaryotic translation.

To assess the physiological relevance of our *in vitro* results, we compared them to ribosome profiling measurements that report A-site occupancy via the ratio of short (21 nt: empty A site) versus long (28 nt: occupied A site) <u>R</u>ibosome-<u>P</u>rotected <u>F</u>ootprint<u>s</u> (RPFs). Prior studies suggested that binding of eRF1 to ribosomes is not rate-limiting for termination, as long RFPs significantly outnumber short RFPs at stop codons (*28*). We confirmed this hypothesis through direct observation of termination sub-steps (**Fig. 6C**), which demonstrated that release factor binding is indeed faster than subsequent ribosomal events. The finding that termination (∼4 s) is fast relative to initiation ∼20 – 60 s (*29, 30*), albeit somewhat slower than elongation (0.05 – 1.4 s per codon (*30*)), suggests the existence of an intricate choreography that prevents the accumulation of ribosomes at stop codons. Consistent with this, ribosomal profiling in eRF1-depleted cells revealed a dramatic increase in queueing of ribosomes at stop codons (*28*).

Eukaryotic termination differs strikingly from the mechanisms described previously for bacterial termination, in which the bacterial namesake of eRF3, RF3, drives departure of eRF1-like factors (RF1/2) from ribosomes (*31*). Instead, eukaryotic termination more closely resembles translation elongation (*5*), in which bacterial EF-Tu and eukaryotic eEF1A assist in the selection of proper tRNAs via a tightly-regulated, multi-step process. In eukaryotic termination, eRF3 (itself an EF-Tu/eEF1A homolog) in complex with GTP quickly delivers eRF1 (a tRNA-shaped protein) to a stop codon in the A site (**Fig. 4A**), similar to the rapid association of eEF1A/tRNA/GTP ternary complex with a sense codon in the A site. eRF3 hydrolyzes GTP to promote its release from the ribosome and facilitate eRF1 accommodation (**Fig. 4D**) (*14*), just as EF-Tu and eEF1A hydrolyze GTP to accelerate their ejection and favor tRNA accommodation (*32, 33*). Furthermore, the energy necessary to eject eRF1 from ribosomes (ΔG^‡^ of 73 ± 34 kJ mol^−1^; **Fig. S6E**) also resembles the energetics of E-site tRNA dissociation (70 kJ mol^−1^; see (*34*)). Thus, the similarities between elongation and eukaryotic termination are not only limited to factor architecture, but also includes the molecular choreography of these processes.

The fidelity of translation elongation is driven in part by kinetic proofreading, in which EF-Tu/eEF1A preferentially rejects non-cognate tRNAs in two sequential steps to boost overall accuracy (*32, 33, 35*). While the basis of termination fidelity is unknown, we consider kinetic proofreading an intriguing model. eRF3 is essential for termination fidelity, as its inclusion boosts specificity by 2,600-fold (*25*). Here we showed eRF3 conformationally unlocks and delivers eRF1 to ribosomes (**Fig. 2** and **4**), and facilitates eRF1 accommodation in an eRF3 GTPase-dependent manner (**Fig. 4**), thus providing eRF3 with multiple opportunities to favor genuine stop codons. Further study of termination sub-step kinetics at near-cognate stop codons will reveal whether proofreading governs eukaryotic termination fidelity.

Mutations that introduce a premature stop codon pose a unique challenge for therapeutic intervention. These mutations trigger premature termination, liberating an incomplete polypeptide, and the defective mRNAs are further degraded via <u>N</u>onsense-<u>M</u>ediated <u>D</u>ecay (NMD) (*36*). To achieve effective therapeutic readthrough of premature stop codons, elongation, termination, and NMD must all be carefully tuned to avoid widespread misregulation of gene expression while still eliciting enough readthrough to alleviate disease. Thus, termination and NMD inhibitors may prove most useful as adjuvants, lengthening the kinetic window for drug-mediated readthrough of premature stop codons. Extension of these single-molecule assays to monitor stop-codon readthrough and NMD will provide the quantitative tools necessary to evaluate combination therapies, paving the way to effective treatments.

## Acknowledgements

We thank Christopher Lapointe, Christine Preston, and Amanda Gilliam Valentic for editing assistance, and also Puglisi and Green lab members for discussion and input.

## Author contributions

MRL, LNL, RG, and JDP conceived the project. MRL, LNL, and JW generated reagents with guidance from RG and JDP. LNL performed and analyzed bulk peptide release experiments with guidance from RG. MRL performed and analyzed all single-molecule experiments (except Fig. S2G, JW) with guidance from JW, AP and JDP. NCC contributed custom single-molecule data analysis scripts. MRL and LNL prepared figures and drafted the initial manuscript, which was revised with input from JW, AP, RG and JDP. All authors read and approved the final manuscript.

## Data and materials availability

Relevant data and expression constructs are available upon request.

## List of Supplementary Materials

Materials and Methods Table S1

Fig S1-S11

References (38 – 54)

## References and Notes

1. L. Frolova et al., A highly conserved eukaryotic protein family possessing properties of polypeptide chain release factor. Nature 372, 701–703 (1994).

2. E. Z. Alkalaeva, A. V. Pisarev, L. Y. Frolova, L. L. Kisselev, T. V. Pestova, In vitro reconstitution of eukaryotic translation reveals cooperativity between release factors eRF1 and eRF3. Cell 125, 1125–1136 (2006).

3. A. Brown, S. Shao, J. Murray, R. S. Hegde, V. Ramakrishnan, Structural basis for stop codon recognition in eukaryotes. Nature 524, 493–496 (2015).

4. C. J. Shoemaker, R. Green, Kinetic analysis reveals the ordered coupling of translation termination and ribosome recycling in yeast. Proc. Natl. Acad. Sci. USA 108, E1392–1398 (2011).

5. D. E. Eyler, K. A. Wehner, R. Green, Eukaryotic release factor 3 is required for multiple turnovers of peptide release catalysis by eukaryotic release factor 1. J. Biol. Chem. 288, 29530–29538 (2013).

6. G. Zhouravleva et al., Termination of translation in eukaryotes is governed by two interacting polypeptide chain release factors, eRF1 and eRF3. EMBO J. 14, 4065–4072 (1995).

7. A. Heuer et al., Structure of the 40S-ABCE1 post-splitting complex in ribosome recycling and translation initiation. Nat. Struct. Mol. Biol. 24, 453–460 (2017).

8. T. Becker et al., Structural basis of highly conserved ribosome recycling in eukaryotes and archaea. Nature 482, 501–506 (2012).

9. K. M. Keeling, X. Xue, G. Gunn, D. M. Bedwell, Therapeutics based on stop codon readthrough. Annu. Rev. Genom. Hum. Genet. 15, 371–394 (2014).

10. M. Mort, D. Ivanov, D. N. Cooper, N. A. Chuzhanova, A meta-analysis of nonsense mutations causing human genetic disease. Hum. Mutat. 29, 1037–1047 (2008).

11. R. Bordeira-Carrico, A. P. Pego, M. Santos, C. Oliveira, Cancer syndromes and therapy by stop-codon readthrough. Trends Mol. Med. 18, 667–678 (2012).

12. M. G. Acker, S. E. Kolitz, S. F. Mitchell, J. S. Nanda, J. R. Lorsch, Reconstitution of yeast translation initiation. Methods Enzymol. 430, 111–145 (2007).

13. J. Wang et al., eIF5B gates the transition from translation initiation to elongation. Nature 573, 605–608 (2019).

14. S. Shao et al., Decoding Mammalian Ribosome-mRNA States by Translational GTPase Complexes. Cell 167, 1229-1240.e1215 (2016).

15. S. Uemura et al., Real-time tRNA transit on single translating ribosomes at codon resolution. Nature 464, 1012–1017 (2010).

16. J. Chen et al., High-throughput platform for real-time monitoring of biological processes by multicolor single-molecule fluorescence. Proc. Natl. Acad. Sci. USA 111, 664–669 (2014).

17. C. Beißel et al., Translation termination depends on the sequential ribosomal entry of eRF1 and eRF3. Nucleic Acids Res. 47, 4798–4813 (2019).

18. J. Flis et al., tRNA Translocation by the Eukaryotic 80S Ribosome and the Impact of GTP Hydrolysis. Cell Rep 25, 2676-2688.e2677 (2018).

19. A. P. Schuller, C. C. Wu, T. E. Dever, A. R. Buskirk, R. Green, eIF5A Functions Globally in Translation Elongation and Termination. Mol. Cell 66, 194-205.e195 (2017).

20. O. Namy, I. Hatin, J. P. Rousset, Impact of the six nucleotides downstream of the stop codon on translation termination. EMBO Rep 2, 787–793 (2001).

21. O. Namy et al., Identification of stop codon readthrough genes in Saccharomyces cerevisiae. Nucleic Acids Res. 31, 2289–2296 (2003).

22. J. R. Wangen, R. Green, Stop codon context influences genome-wide stimulation of termination codon readthrough by aminoglycosides. eLife 9, e52611 (2020).

23. E. M. Youngman, L. Cochella, J. L. Brunelle, S. He, R. Green, Two distinct conformations of the conserved RNA-rich decoding center of the small ribosomal subunit are recognized by tRNAs and release factors. Cold Spring Harb Symp Quant Biol 71, 545–549 (2006).

24. D. E. Eyler, R. Green, Distinct response of yeast ribosomes to a miscoding event during translation. RNA 17, 925–932 (2011).

25. G. Indrisiunaite, Doctoral thesis, comprehensive summary, Acta Universitatis Upsaliensis, Uppsala (2019).

26. H. Song et al., The crystal structure of human eukaryotic release factor eRF1--mechanism of stop codon recognition and peptidyl-tRNA hydrolysis. Cell 100, 311–321 (2000).

27. Z. Cheng et al., Structural insights into eRF3 and stop codon recognition by eRF1. Genes Dev. 23, 1106–1118 (2009).

28. C. C. Wu, B. Zinshteyn, K. A. Wehner, R. Green, High-Resolution Ribosome Profiling Defines Discrete Ribosome Elongation States and Translational Regulation during Cellular Stress. Mol. Cell 73, 959-970.e955 (2019).

29. P. Shah, Y. Ding, M. Niemczyk, G. Kudla, J. B. Plotkin, Rate-limiting steps in yeast protein translation. Cell 153, 1589–1601 (2013).

30. D. Chu et al., Translation elongation can control translation initiation on eukaryotic mRNAs. EMBO J. 33, 21–34 (2014).

31. D. V. Freistroffer, M. Y. Pavlov, J. MacDougall, R. H. Buckingham, M. Ehrenberg, Release factor RF3 in E.coli accelerates the dissociation of release factors RF1 and RF2 from the ribosome in a GTP-dependent manner. EMBO J. 16, 4126–4133 (1997).

32. T. Pape, W. Wintermeyer, M. V. Rodnina, Complete kinetic mechanism of elongation factor Tu-dependent binding of aminoacyl-tRNA to the A site of the E. coli ribosome. EMBO J. 17, 7490–7497 (1998).

33. T. Pape, W. Wintermeyer, M. Rodnina, Induced fit in initial selection and proofreading of aminoacyl-tRNA on the ribosome. EMBO J. 18, 3800–3807 (1999).

34. J. Choi, J. D. Puglisi, Three tRNAs on the ribosome slow translation elongation. Proc. Natl. Acad. Sci. USA 114, 13691 (2017).

35. A. B. Loveland, G. Demo, A. A. Korostelev, Cryo-EM of elongating ribosome with EF-Tu•GTP elucidates tRNA proofreading. Nature 584, 640–645 (2020).

36. F. He, A. Jacobson, Upf1p, Nmd2p, and Upf3p regulate the decapping and exonucleolytic degradation of both nonsense-containing mRNAs and wild-type mRNAs. Mol. Cell. Biol. 21, 1515–1530 (2001).

37. E. Svidritskiy, A. F. Brilot, C. S. Koh, N. Grigorieff, A. A. Korostelev, Structures of yeast 80S ribosome-tRNA complexes in the rotated and nonrotated conformations. Structure 22, 1210–1218 (2014).

