## Supplementary Materials for "Mechanisms that ensure speed and fidelity in eukaryotic translation termination"

**Materials and Methods**

Cloning, expression, purification, and fluorescent labeling of eukaryotic release factors:

Full-length eRF1 and eRF3∆165N from *S. cerevisiae*, PCR-amplified from pTYB2 expression vectors (*5*), were inserted into 1C expression vectors (UC Berkeley MacroLab, 6xHis-MBP-TEV-ORF), either with or without an ybbR tag (DSLEFIASKLA) inserted between the TEV and ORF regions, using standard UC Berkeley Macrolab ligation-independent cloning protocols. Expression vectors were transformed individually into *Escherichia coli* Rosetta2 (DE3) pLysS (Novagen), and overexpressed separately in Terrific Broth media (Fisher) by induction at an OD600 of 0.5-0.6 with 1 mM IPTG at 16˚ C. For each purification, 3L of cells were harvested by centrifugation at 4,000 x g after 16 hours, resuspended in Ni-A buffer (500 mM NaCl, 25 mM Tris, 30 mM Imidazole, 10% Glycerol, 0.2 mM TCEP, 1 mM PMSF, 1 µg/mL Leupeptin, 2 µg/mL Pepstatin A, pH 7.5 at room temperature) supplemented with a cOmplete Mini EDTA-free protease inhibitor cocktail tablet (Roche) and either 0.5 mM EDTA (eRF1) or 100 µM GTP (eRF3∆165N), flash frozen in liquid nitrogen, and stored at -80 ˚C.

Cells were lysed by sonication (MISONIX, 20 s on / 40 s off, 6 min total on time), cellular debris pelleted by centrifugation in an SS34 rotor at 18,000 rpm, and the supernatant loaded onto a 5 mL HisTrap HP column (GE) using a peristaltic pump at roughly 2 mL/min flow rate. Columns were washed with 50 CV of Ni-A, 25 CV of High Salt Ni-A (same as Ni-A, but with 1M NaCl) and finally 10 CV of Ni-A, with each of these buffers supplemented with either 0.5 mM EDTA (eRF1) or 100 µM GTP (eRF3∆165N). Release factors were eluted in Ni-B (same as Ni-A, but with 300 mM Imidazole and no EDTA/GTP), fractions collected, combined with 2 mg of TEV protease (UC Berkeley Macrolab), injected into a Slide-A-Lyzer dialysis cassette (10K MWCO, 15 mL, Thermo Scientific), and dialyzed into 1L of Ni-A buffer overnight on a stir plate. TEV-cleaved release factors were run through HisTrap HP column using a peristaltic pump (to separate away TEV and uncleaved release factors), diluted 1:10 with low salt buffer (5 mM NaCl, 5 mM HEPES pH 7.5), and loaded onto a 1 mL Q FF column (GE) using a peristaltic pump. After a further 20 CV wash with 1:10 Ni-A:Low salt buffer, factors were eluted using Ni-A into a ~2 mL volume, flash frozen in liquid nitrogen as 0.5 mL aliquots, and stored at -80 ˚C. Later (after labeling and free dye separation, in the case of labeled factors), factors were thawed on ice and run over a 320 mL Superdex 200 column pre-equilibrated in sizing buffer (200 mM KCl, 20 mM HEPES, 1.5 mM MgCl_2_, 10% Glycerol, 2 mM DTT, pH 7.5). Fractions containing intact release factors, as judged by SDS-PAGE and Coomassie staining, were pooled, concentrated to ~10 mg/mL per A_280_ readings (ε = 36,330 M^-1^ cm^-1^ for eRF1, 44,350 M^-1^ cm^-1^ for eRF3∆165N), aliquoted, flash frozen in liquid nitrogen, and stored at -80 ˚C. Purifications typically yielded milligram quantities of release factors.

CoA-conjugated Cyanine dyes for labeling were obtained as described previously (*13*). Prior to labeling, eRF1 or eRF3∆165N were dialyzed overnight at 4 ˚C into labeling buffer (100 mM NaCl, 50 mM HEPES KOH pH 7.5, 10 mM MgCl2, 5 mM DTT) using Slide-A-Lyzer MINI dialysis cassettes (10K MWCO, 0.5 mL) with ≥ 1,000-fold excess of buffer. In the morning, factors were dialyzed again into a fresh ≥ 1,000-fold excess of labeling buffer for at least 2 h, and concentration measured by A_280_. Labeling reactions were assembled as follows (0.5 – 2 mL final volume), wrapped in foil, and allowed to incubate at 4˚ C overnight for at least 20 h: 25 µM eRF1-N_ybbR or eRF3∆165N-N_ybbR, 50 µM CoA-conjugated dye, 4 µM Sfp (purified as described, (*38*)), and 100 mM DTT. Next, labeling reactions were applied to 10DG desalting columns (BioRad) pre-equilibrated in sizing buffer, with one 10DG column for every 0.5 mL of labeling reaction. Factors were loaded and eluted using the BioRad 10DG minimum dilution protocol, and 1 mL fractions collected. Fractions that contained labeled release factors yet no free dye (selected by SDS-PAGE separation and fluorescence scans with a Typhoon imager) were pooled, further purified using size exclusion chromatography, and stored as described above. Labeling efficiencies ranged from 45-80%.

Bulk elongation and termination reactions:

Translation initiation factors eIF1, eIF1A, eIF5, and eIF5B were expressed and purified from *E. coli*, and eIF2 was expressed and purified from *S. cerevisiae* as previously described (*12, 24*). The translation elongation factors eEF2 and eEF3 were purified from S. cerevisiae as previously described (*19*). The translation elongation factor eIF5A was purified from E. coli as previously described (*19, 39*). The translation termination factors eRF1 (and the inactive eRF1 mutant G180A) were purified from *E. coli* as previously described (*40*), and eRF3 was purified from *E. coli* as previously described (*5*).

80S initiation complexes were formed as previously described (*19, 41*). Non-acylated initiator and charged lysine tRNAs were purchased from tRNA probes (College Station, TX). Phenylalanine tRNA was purchased from Sigma, and charged using purified synthetase; and tRNA^Met^_i_ was charged with EasyTag Methionine L‐[35S] (Perkin Elmer) and purified synthetase as previously described (*41*). mRNAs were purchased from IDT. Initiation complexes were tested for reactivity with Puromycin as previously described and used to normalize initiation complex activity (*41*).

Elongation reactions were performed as previously described with minor adjustments to elongate to the stop codon and be immediately followed by translation termination to quantify peptide release (*19, 41*). Briefly, aa‐tRNA ternary complex was formed by incubating aa‐tRNA (1.5–2 μM), eEF1A (15 μM), and 1 mM GTP, in 1× Buffer E for 10 min at 26° C. Limited amounts of 80S initiation complexes (2-3 nM) were then mixed with aa‐tRNA ternary complex (50-150 nM), eEF2 (1.5 µM), eEF3 (4 μM), eIF5A (1.25 μM), ATP (3 mM), and GTP (2 mM). Reactions were incubated at 26°C for 5 to 10 minutes. While elongation mixture incubated, eRF1 (6 µM) and eRF3 (10 μM) were mixed with 1 X Buffer E (2.5 mM Mg(OAc)_2_) and 1mM GTP/Mg(OAc)_2_ and any additional factors (6 µM) on ice (termination mix). Elongation reaction mixture (6 µL) was then mixed with the termination mixture (3 µL) and incubated at 26° C (unless stated otherwise). Final concentrations in termination reactions were as follows: 2 µM eRF1, 3.3 µM eRF3, 1.5 nM ribosome complexes, 1.3 µM eEF1A, 1 µM eEF2, 2.7 µM eEF3, and 833 nM eIF5A. Time points were removed and quenched into an equal volume of 10% formic acid. Samples (1 µL) were spotted on TLC cellulose plates (3.5 cm from bottom, 0.9 cm apart; EMD Millipore, 1055770001). TLC plates were equilibrated with pyridine acetate buffer (5 ml pyridine, 200 ml acetic acid in 1 L, pH 2.8) before electrophoresis at 1400 V for 25 min. TLC plates were developed using a Typhoon FLA 9500 Phosphorimager system (GE Healthcare Life Sciences) and quantified using ImageQuantTL (GE Healthcare Life Sciences). Time courses were fit to single‐exponential kinetics using Prism version 8 (GraphPad Software Inc.).

Reconstitution for single-molecule assays:

Yeast initiation factors and ribosomal subunits were purified as described previously (*13*). 80S complexes for single-molecule assays were generated using an adapted version of published protocols (*13, 42*). All samples were prepared in 1X Rec buffer (30 mM HEPES-KOH pH 7.5, 100 mM KOAc, 3 mM Mg(OAc)_2_). To first generate 3X eIF2•Met-tRNA^Met^_i_•GTP, 3.9 µM eIF2 and 1 mM GTP/Mg(OAc)_2_ were incubated together at 30 ˚C for 10 min, combined with Met-tRNA^Met^_i_ (2.7 µM) and again incubated at 30 ˚C for 5 min. 10X eIF1•eIF1A•GTP was generated by mixing 10 µM eIF1, 10 µM eIF1A, and 1 mM GTP-Mg(OAc)_2_ together on ice. 48S Pre-Initiation Complexes (48S PIC) was generated by incubating a mixture of eIF2•Met-tRNA^Met^_i_•GTP (1X final), eIF1•eIF1A•GTP (1X final), 1 mM GTP/Mg(OAc)_2_, mRNA (3 µM, purchased from IDT), and 40S (80 nM) together for 15 min at 30 ˚C. A 60S mixture was assembled separately by mixing 1 µM eIF5, 2 µM eIF5B∆396N, and 60S (either 200 nM unlabeled or 100 nM labeled) together on ice. 48S PIC and 60S mixtures were then combined, incubated at room temperature for 10 min, and supplemented with casein (62.5 µg/mL final concentration, added as a blocking agent).

RS experiments and data processing:

Real-time single-molecule measurements in ZMWs were achieved using an RS II (Pacific Biosciences) customized as described previously (*16*). ZMW chips were prepared as described previously (*13*), except pre-formed 80S complexes were immobilized here rather than 48S. 20 µL of imaging buffer was added to the chip with 1X Rec buffer, oxygen scavenging system (4 mM PCA, 4 mM TSY, 2X PCD (protocatechuate-3,4-dioxygenase, added last); all purchased from Pacific Biosciences), 62.5 µg/mL casein, and 1 mM GTP-Mg(OAc)_2_ (or analogs) and/or 1 mM ATP-Mg(OAc)_2_ matching conditions of a given experiment as appropriate. After the commencement of real-time data acquisition, 20 µL of a “delivery mix” was pipetted onto the chip, resulting in a 2-fold dilution of factors; concentrations referenced throughout the main text account for this dilution.

For all experiments, a 10X release factor mix was first generated by mixing (final concentrations listed) 1 mM GTP-Mg(OAc)_2_ or 1 mM GTPγS-Mg(OAc)_2_ (GTP purchased from Sigma, GTPγS from Jena Biosciences), eRF1, eRF3, and sizing buffer (200 mM KCl, 20 mM HEPES, 1.5 mM MgCl_2_, 10% Glycerol, 2 mM DTT, pH 7.5; added such that the volume of eRF1, eRF3 and sizing buffer was 60% of the total delivery mix) and 0.2X Rec buffer together on ice. 10X release factor mix for RS experiments with unlabeled eRF3 (eRF1 dynamics, **Fig. 2C-D**; Met-tRNA^i^-eRF1 FRET and termination assays, **Fig. 5-6**) contained 10 µM eRF3∆165N. Delivery mixes for all RS experiments contained (final concentrations) 1X Rec buffer, 1 mM GTP-Mg(OAc)_2_, 1X release factor mix, 62.5 µg/mL casein, and oxygen scavenging system (see above).

Tracking of eRF1 dynamics (**Fig. 2** and **S6F**) was conducted with 532-nm laser illumination at 0.32 µW/µM^2^ (unless otherwise stated), with 10 frames collected per second for 10 minutes at 20 ˚C. For the eIF5A-containing experiment described in **Fig. S6**, delivery mix was supplemented with 2 µM eIF5A, and imaging conducted instead at 30 ˚C. Tracking of eRF3 dynamics (**Fig. 3**) was conducted under dual-illumination (532-nm laser at 0.32 µW/µM^2^, 642-nm laser at 0.10 µW/µM^2^), with 10 frames collected per second for 10 minutes at 20 ˚C. For co-tracking of eRF1/eRF3 dynamics (**Fig. 4** and **S9F**), imaging buffer and delivery mixes were further supplemented with 1 µM double-stranded DNA oligonucleotide and 5 mg/mL BSA (as blocking agents, described previously (*16*)), and imaging was conducted at 532-nm laser illumination at 0.32 µW/µM^2^, 10 frames per second, for 10 minutes at 20 ˚C. For the apo experiment listed in **Fig. S4A** and **S4E**, GTP-Mg(OAc)_2_ was omitted from 10X release factor mix, delivery mix, and 10X release factor mixes. Paromomycin was included in the delivery mix at 100 µM final concentration (to account for 2-fold dilution after addition to the chip) for the drug-containing experiment in **Fig. S9F**, and imaging conducted instead at 30 ˚C.

For tracking of Cy3-labeled Met-tRNA^i^ / Cy5-labeled G180A-eRF1 FRET (**Fig. 5A-B**), 80S complexes were prepared as described above, except 190 nM Cy3-labeled Met-tRNA^i^ (acquired from tRNA Probes) was instead used in the eIF2•Met-tRNA^Met^_i_•GTP mix. Where listed, 4 mM puromycin was added to the delivery mix. Imaging was conducted at 532-nm laser illumination at 0.32 µW/µM^2^, 10 frames per second, for 15 minutes at 20 ˚C. For real-time monitoring of termination (**Fig. 5** and **6**), delivery mixes were further supplemented with an 8X Cy3-labeled (or Cy3.5-labeled, when noted) Phe-tRNA^Phe^ ternary complex (100 nM final in delivery mix, prepared as described previously, (*13*)), 1 µM eEF2, 0.4 µM eEF3, 2 µM eIF5A, and 1 mM ATP-Mg(OAc)_2_. Imaging was conducted with 532-nm laser illumination at 0.32 µW/µM^2^ (unless otherwise stated), with either 8 or 10 frames collected per second, for 15 minutes at 30 ˚C (unless otherwise stated).

For all experiments, data were processed using a suite of custom-written Matlab scripts (described previously (*16, 43*), and available upon request). After filtering all traces for expected FRET events, traces were manually inspected for expected fluorescence events of a particular experiment. For Cy3 eRF1/Cy5-60S FRET (**Fig. 2**), only traces that displayed photobleaching of the 60S subunit (and a simultaneous increase in Cy3 fluorescence) were selected for analysis. In all cases, association time was marked as the time window between deposition of the delivery mix on the surface (which typically varied from ~5-15 s, but was recorded for every individual experiment) and the appearance of expected fluorescence. Single- and double-exponential function fits were achieved as described previously (*16*), and pseudo-first order rate constants obtained via linear regression model fitting to apparent rate constants in Prism 8 (GraphPad Software Inc.), with reported error ranges denoting 95% CIs. Probability distributions (plots of Probability vs. Time) were rendered in Prism 8. FRET intensities were fit to single gaussian models and plots rendered in Prism 8, with error ranges denoting 95% CI of mean FRET. For single-molecule termination assays (**Fig.** **5** and **6**), only traces that displayed at least three frames medium-FRET states were included for analysis (unless otherwise noted, such as lifetime analysis in **Fig. S5J**), as short-lived transitions from high- to medium-FRET are difficult to distinguish from a mid-frame photobleaching event. Pre-termination state lifetimes were fit using the cftool function in Matlab the following two-step, irreversible kinetic model (described previously (*44*)):

$$C\left( t \right)=1- \frac{b}{b-a}e^{-at}-\frac{a}{a-b}e^{-bt}$$

Pre-termination state lifetimes were calculated as the sum of 1/*a* and 1/*b* (referred to as τ_A_ and τ_B_, respectively). 95% CIs yielded for rates a and b were similarly used to determine error ranges in seconds for upper and lower bounds (deviation from τ_A_ and τ_B_ noted as σ_A,upper_, σ_A,lower_, σ_B,upper_, and σ_B,lower_). The upper bound of the 95% CI was calculated as: σ_AB,upper_ = ((σ_A,upper_)^2^ + (σ_B,upper_)^2^)^0.5^, and lower bound of the 95% CI was calculated as: σ_AB,lower_ = ((σ_A,lower_)^2^ + (σ_B,lower_)^2^)^0.5^. Post-termination state lifetimes were fit to a single-exponential function.

Co-arrival likelihood calculations:

We used the same calculations described previously (*45*) to assess the likelihood of sequential eRF1 and eRF3 binding occurring in the same frame by random chance. When considering any particular eRF1 binding event, the probability that eRF3 would also appear to bind at the same time is:

$$p=1- e^{-k_{obs}*t*n}$$

Where *t* is the duration of each frame, *n* is number of frames, and *k_obs_* is the eRF3 association rate. Here, *t* = 0.1 s, *n* = 1.5 (to also account for the possibility of eRF3 binding in the latter half of the preceding frame), and *k_obs_* = 0.006268 (from **Fig. S3**), yielding *p* = 0.000940. Analysis of the co-arrival datasets depicted in **Fig. 4C**, accounting for all eRF1 binding events observed irrespective of eRF3, yields only 1.8 – 2.2 co-arrival events expected by chance (25 nM, 1,968 eRF1 binding events; 50 nM, 1,545; 100 nM, 2,384), far fewer than the 122 – 192 co-arrival events observed in these experiments (192, 122, and 153 respectively; **Fig. S4C**). Notably, our ability to observe eRF1/eRF3 co-arrival here is also hampered by incomplete labeling of both eRF1 (46%) and eRF3 (74%).

TIRF experiments and data processing:

TIRF imaging was conducted on a home-built, prism-based instrument described previously (*46, 47*). 80S ribosomes were immobilized on neutravidin-derivatized quartz slides, and unbound ribosomes washed away as described previously (*13*). Immediately prior to data acquisition, a mixture of termination factors was added to the surface that contained 1 mM GTP-Mg(OAc)_2_, 10 nM Cy-labeled eRF1, 1 µM eRF3∆165N, and oxygen scavenging system (described above) all in 1X Rec buffer (prepared as described above). A 532-nm laser at 1 kW cm^-2^ was used for illumination, and movies collected using the MetaMorph software package (Molecular Devices) at a rate of 10 frames per second for a duration of 1 to 5 minutes. Movies were processed and traces selected using in-house MATLAB scripts described previously (available on Github, (*43*)) and Spartan (*48*). FRET states were assigned using vbFRET (*49*), and FRET efficiency histograms rendered in Prism 8 (GraphPad Software Inc.).

Eyring analysis: Following work described by others (*34, 50, 51*), pre- and post-termination kinetics (k_obs_) and relevant temperatures (*T*) were fit first via linear regression to the Eyring equation:

$$\ln\left( k_{obs} \right)= -\frac{\Delta H^{\ddagger}}{R}\cdot\frac{1}{T}+\ln\left( \frac{k_{B}T}{h} \right)+\frac{\Delta S^{\ddagger}}{R}$$

Where h is the plank constant, kB is the Boltzman constant, and R the ideal gas constant. Next, the Gibbs Free Energy of the transition state was calculated as follows:

$$\Delta G^{\ddagger}=\Delta H^{\ddagger}-T\Delta S^{\ddagger}$$

Error ranges noted for ∆G^‡^, ∆H^‡^, and ∆S^‡^ represent standard error.

Sequence alignments: Release factor sequences were aligned using the T-Coffee webserver (*52*) and visualized using Boxshade.

**Table S1:**

| mRNA | Sequence (**ORF in bold**) |
| --- | --- |
| M-F-K-K-Stop | GAA UCU CUC UCU CUC UCU **AUG UUC AAA AAA UAA** CUC UCU CUC UCU CUC |
| M-Stop (UAA) | /5BiosG/GAAUCUCUCUCUCUCUCUCUCUCUCUCUCUCUCUCUCUCUCU**AUGUAA**AAA |
| M-F-Stop (UAA) | /5BiosG/GAAUCUCUCUCUCUCUCUCUCUCUCUCUCUCUCUCUCUCUCU**AUGUUCUAA**AAA |
| M-F-Stop (UAG) | /5BiosG/GAAUCUCUCUCUCUCUCUCUCUCUCUCUCUCUCUCUCUCUC**AUGUUCUAG**AAA |
| M-F-Stop (UGA) | /5BiosG/GAAUCUCUCUCUCUCUCUCUCUCUCUCUCUCUCUCUCUCUCU**AUGUUCUGA**AAA |
| M-F-Stop (UGAC) | /5BiosG/GAAUCUCUCUCUCUCUCUCUCUCUCUCUCUCUCUCUCUCUCU**AUGUUCUGA**CAA |
| M-F-Stop (UAU) | /5BiosG/GAAUCUCUCUCUCUCUCUCUCUCUCUCUCUCUCUCUCUCUCU**AUGUUCUAU**AAA |
| M-F-Stop (Str. Readthrough) | /5BiosG/GAAUCUCUCUCUCUCUCUCUCUCUCUCUCUCUCUCUCUCUCU**AUGUUCUAG**CAAUUAAAA |
| M-F-Stop (Med. Readthrough) | /5BiosG/GAAUCUCUCUCUCUCUCUCUCUCUCUCUCUCUCUCUCUCUCU**AUGUUCUAG**CAAUCAAAA |
| M-F-Stop (Weak Readthrough) | /5BiosG/GAAUCUCUCUCUCUCUCUCUCUCUCUCUCUCUCUCUCUCUCU**AUGUUCUAG**CAAAGAAAA |

**Fig. S1-S11:**

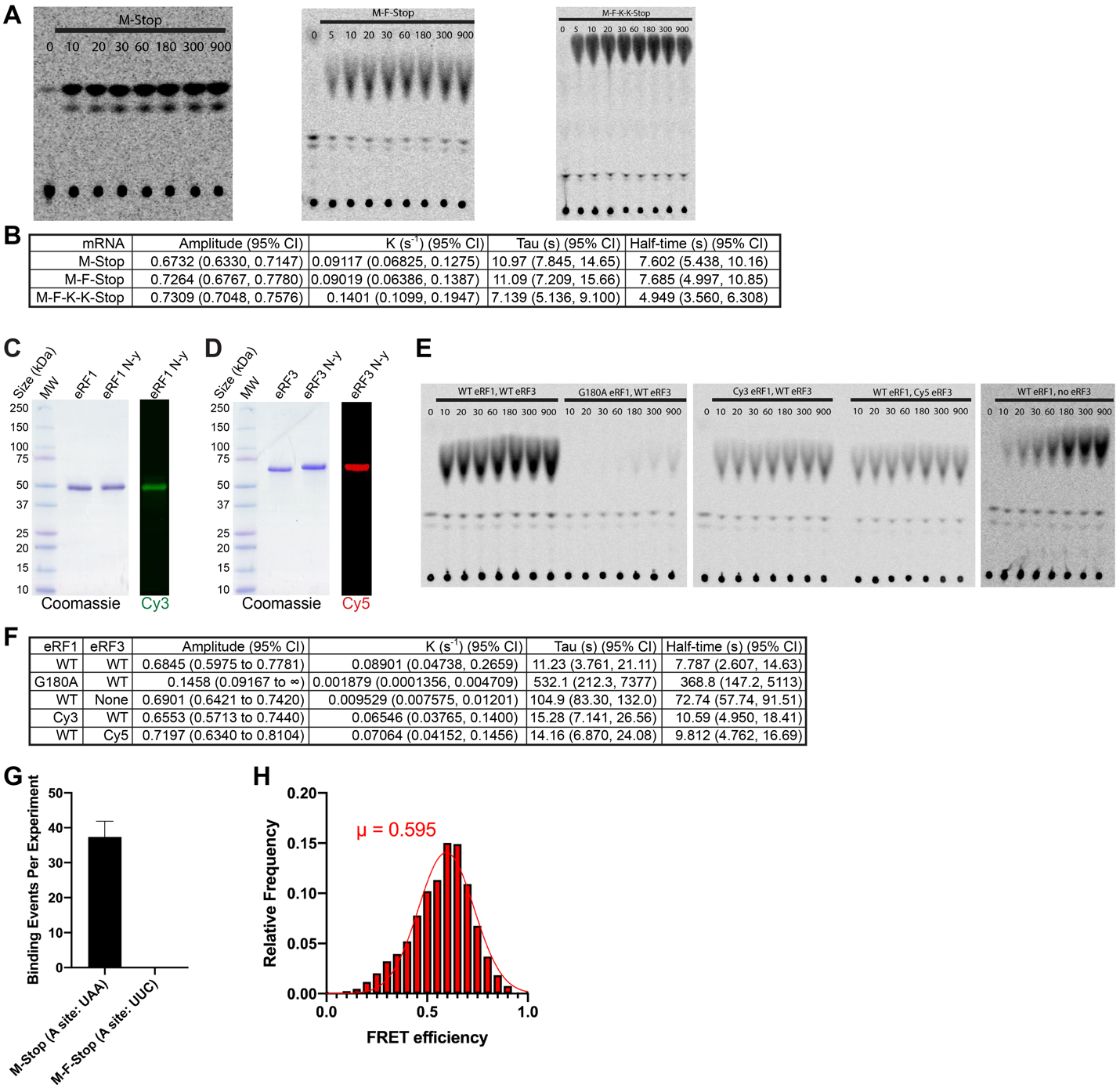

**Fig. S1**: Tools for single-molecule studies of translation termination. (**A-B**) Peptides are released at similar rates from model mRNAs, (**A**) example TLCs (time in seconds) and (**B**) fitting statistics. (**C-D**) Purification of wild-type and fluorescently-labeled (**C**) eRF1 and (**D**) eRF3∆165N (“eRF3”). Labeling achieved via attachment of CoA-conjugated Cy dyes using Sfp chemistry (*38*). (**E-F**) Fluorescent labeling of eRF1 and eRF3 does not impair their ability to liberate peptides from M-F-Stop ribosomes, (**E**) example TLCs (time in seconds) and (**F**) fitting statistics. (**G**) Binding events observed per TIRF experiment with M-Stop or M-F-Stop ribosomes. (**H**) eRF1/60S FRET efficiency observed in TIRF with M-Stop ribosomes.

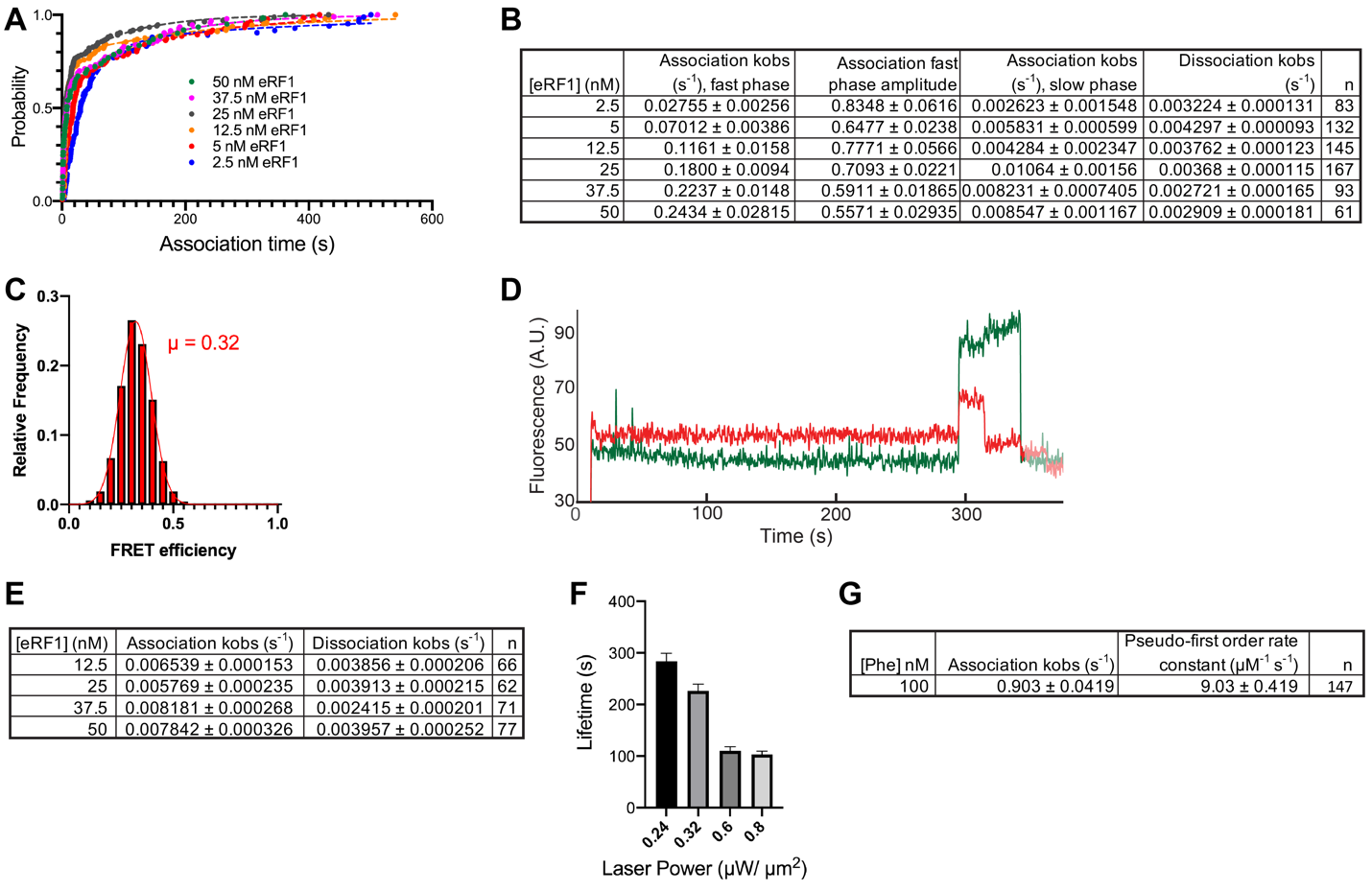

**Fig. S2**: Ribosomal dynamics of eRF1 in the presence or absence of eRF3. (**A-B**) Concentration-dependent binding of eRF1 observed in the presence of 500 nM eRF3, (**A**) 0-600 s, and (**B**) statistics on fits. (**C**) FRET observed between Cy3-eRF1 and Cy5-L5-60S. (**D**) Example of slow eRF1 binding observed without eRF3. (**E**) Concentration-independent binding of eRF1 observed without eRF3, statistics on fits. (**F**) eRF1 lifetime observed at different 532-nm laser powers. (**G**) Statistics on fit of Phe-tRNA^Phe^ ternary complex binding to its cognate UUC A-site codon.

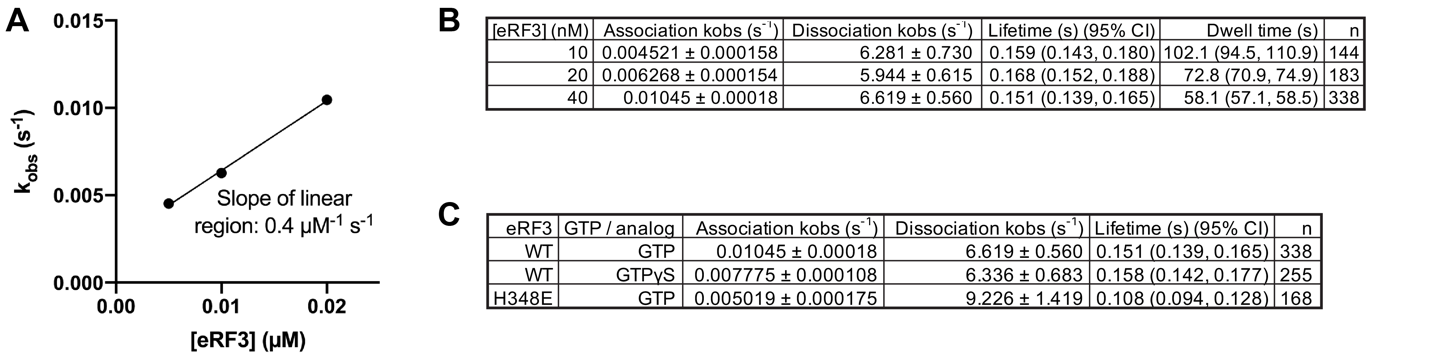

**Fig. S3**: Ribosomal dynamics of eRF3. (**A-B**) Observed rates of eRF3 binding to M-stop ribosomes (k_obs_), (**A**) plotted vs. eRF3 concentration and (**B**) statistics on fits. (**C**) Statistics on fits to observed eRF3 GTPase-dependent M-Stop ribosomal dynamics.

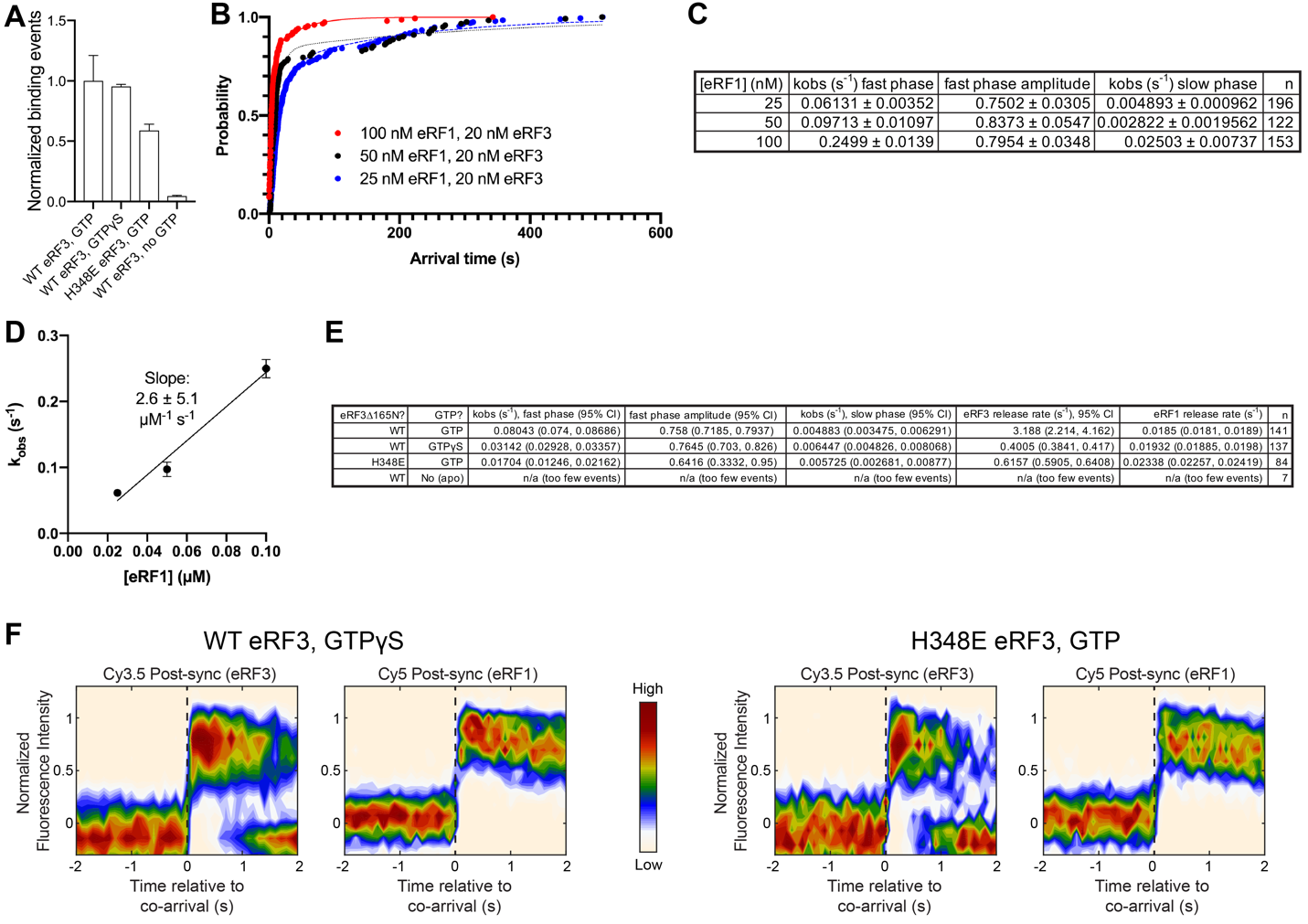

**Fig. S4**: Simultaneous tracking of eRF1 and eRF3 dynamics. (**A**) Observed ternary complex binding events, with GTP/analogs or a GTPase-defective eRF3 mutant. (**B**) Probability distribution for association of ternary complex with M-stop ribosomes, 0-600 s. (**C-D**) Ternary complex arrival rates, (**C**) statistics on fits and (**D**) plotted vs. different concentrations of eRF1. (**E**) Statistics on fits for release factor dynamics, with GTP/analogs or a GTPase-defective eRF3 mutant. (**F**) Post-synchronization plot of fluorescence changes observed upon simultaneous binding of eRF1 and eRF3 (denoted as a dashed, black vertical line), either with wild-type eRF3 and GTPγS (*left*) or H348E eRF3 and GTP (*right*).

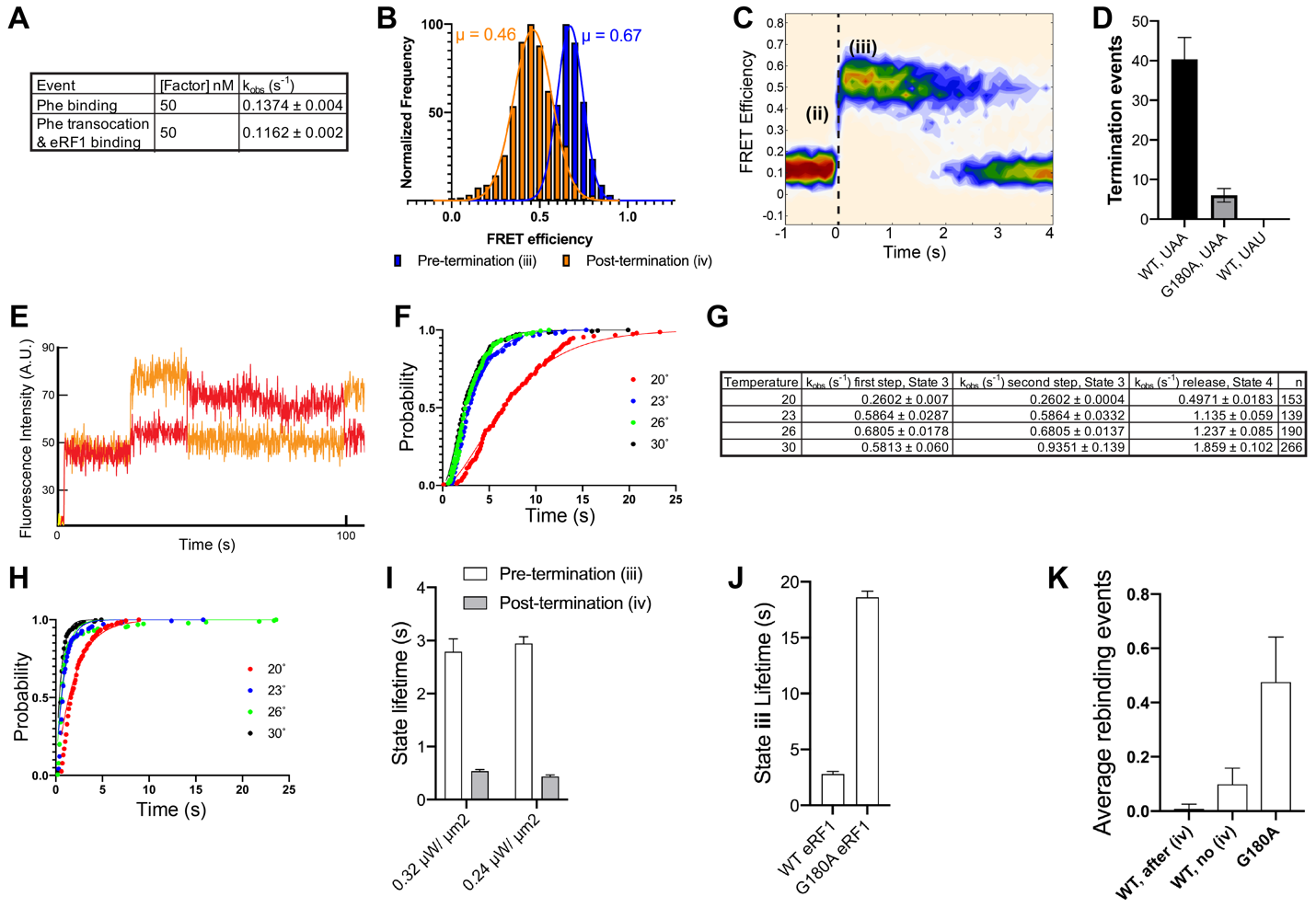

**Fig. S5**: Monitoring peptidyl-tRNA bond status by FRET. (**A**) Observed rates (k_obs_) of Phe tRNA binding (i) or Phe translocation and eRF1 binding (ii). (**B**) FRET observed between P-site Cy3-Phe tRNA and eRF1. (**C**) Cy3-Phe/Cy5-eRF1 FRET, post-synchronized to end of state ii. (**D**) Termination events observed with wild-type or G180A eRF1 at a stop (UAA) or near-cognate stop (UAU), 30 ˚C. (**E**) Example of FRET observed with G180A eRF1, 30˚ C. (**F**) Pre-termination state lifetime and fits at different temperatures. (**G**) Pre- and post-termination state lifetime fitting statistics. (**H**) Post-termination state lifetime fits at different temperatures. (**I**) Pre- and post-termination state lifetimes at different laser powers. (**J**) State iii lifetime observed with wild-type or G180A eRF1, 30 ˚C. (**K**) Frequency of eRF1 rebinding events observed with wild-type eRF1 to either post-termination ribosomes (“after (iv)”), or pre-termination ribosomes (“no (iv)”), and G180A eRF1 to pre-termination ribosomes, 30 ˚C.

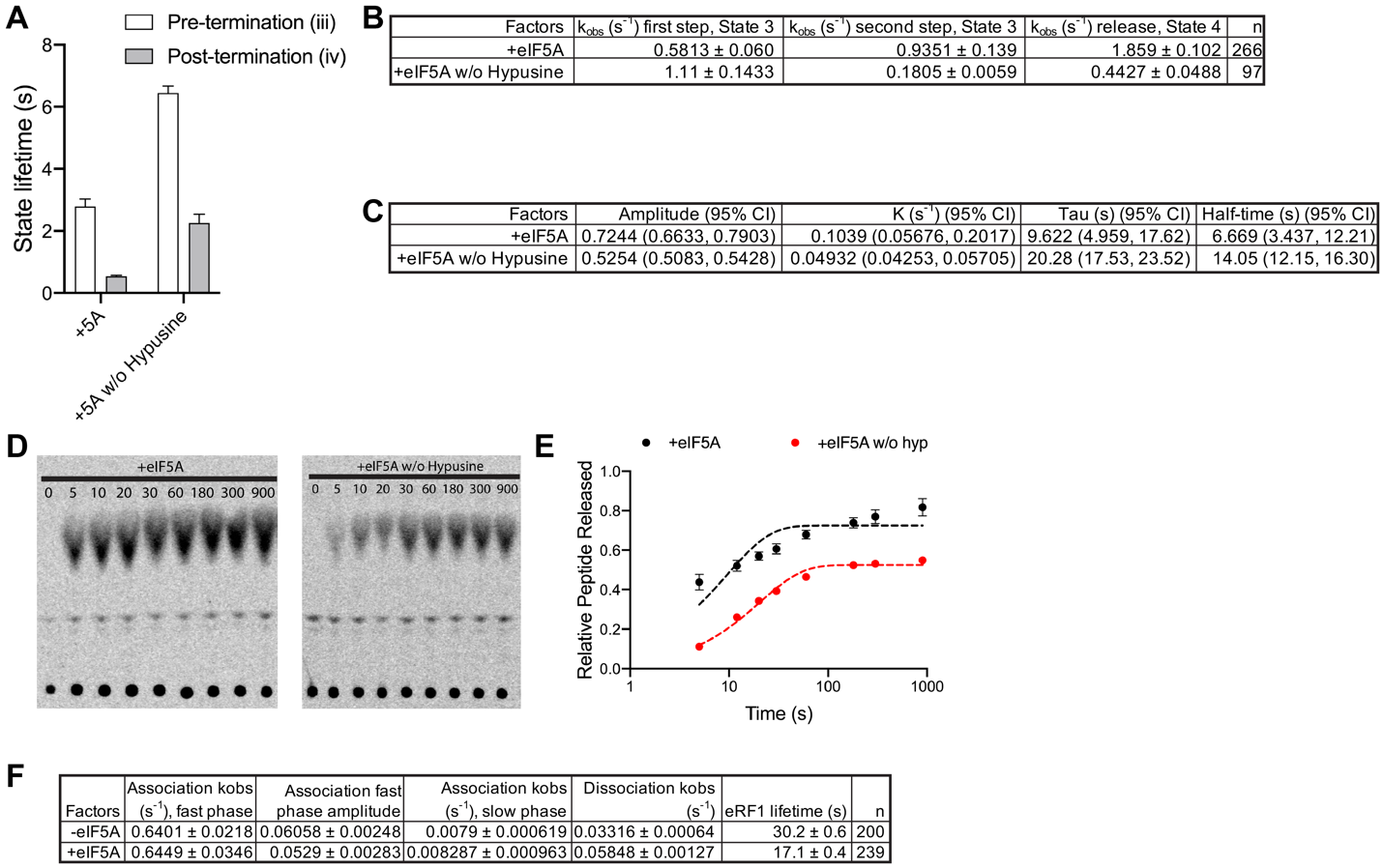

**Fig. S6**: eIF5A promotes peptidyl-tRNA bond hydrolysis and eRF1 release. (**A-B**) Substitution of eIF5A with an inactive variant (lacking its critical hypusine modification (*53, 54*)) significantly lengthened the pre- and post-termination states lifetimes. (**A**) State lifetimes and fits and (**B**) fitting statistics. (**C-E**) Impact of eIF5A peptide release rates with M-F-Stop ribosomes. (**C**) fitting statistics, (**D**) example TLCs (time in seconds), and (**E**) fraction of peptide released with one-phase association fits. (**F**) eIF5A accelerates release of Cy5-labeled eRF1 from Cy3-60S-labeled M-Stop ribosomes at 30 ˚C. The remaining discrepancy between eRF1 lifetime in single-molecule termination experiments (**Fig. 5**, 3-4 s on ribosome) vs. that seen here with eIF5A (17.1 s) may be attributable to the technical difficulties of reliably identifying short-lived, low-FRET events (thus skewing the eRF1/60S FRET distribution towards longer events), and/or because the events we focused on in **Fig. 5** all result in termination, which favors eRF1 release.

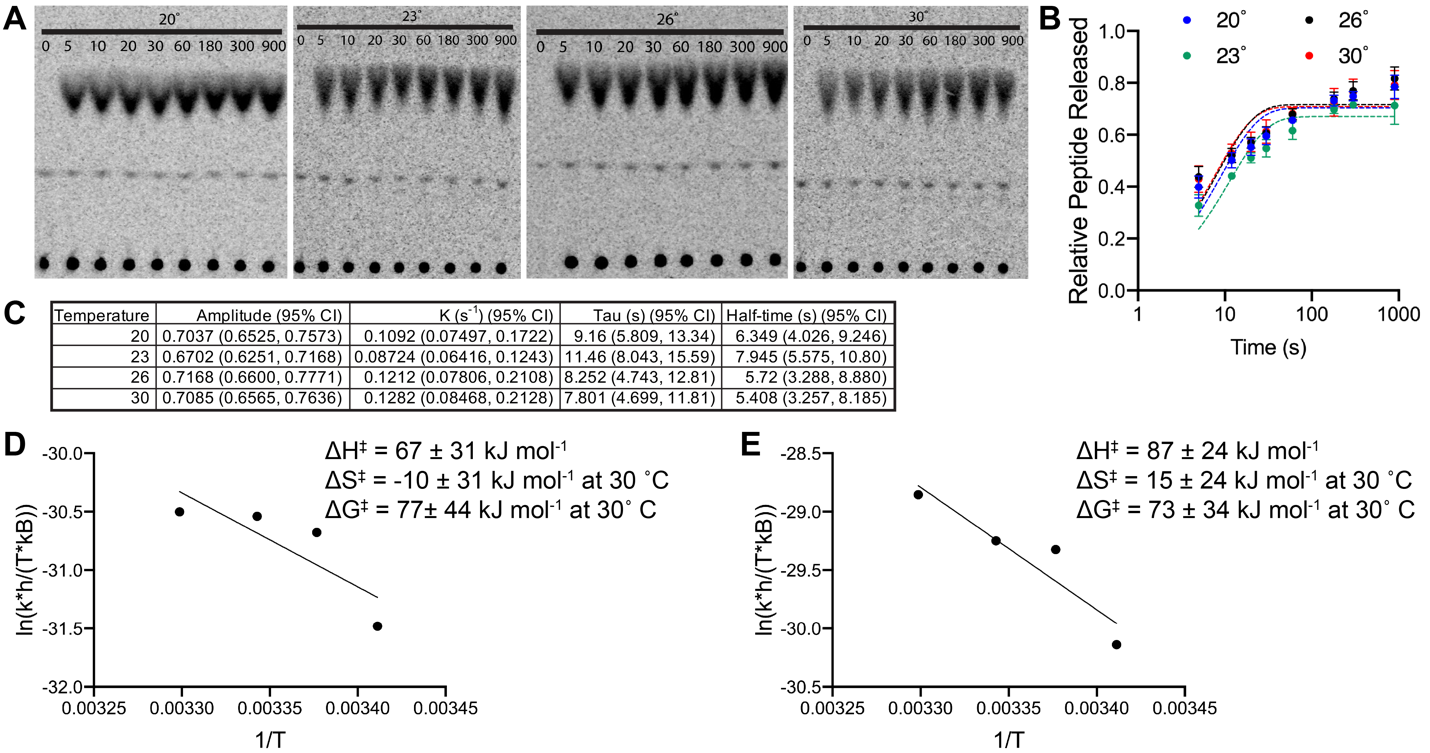

**Fig. S7:** The impact of temperature on termination kinetics. (**A-C**) Rate of peptide release from M-F-Stop ribosomes increases with temperature. (**A**) example TLCs (time in seconds), (**B**) fraction of peptide released with one-phase association fits, and (**C**) fitting statistics. (**D-E**) Eyring plots of (**D**) pre- and (**E**) post-termination rates at different temperatures.

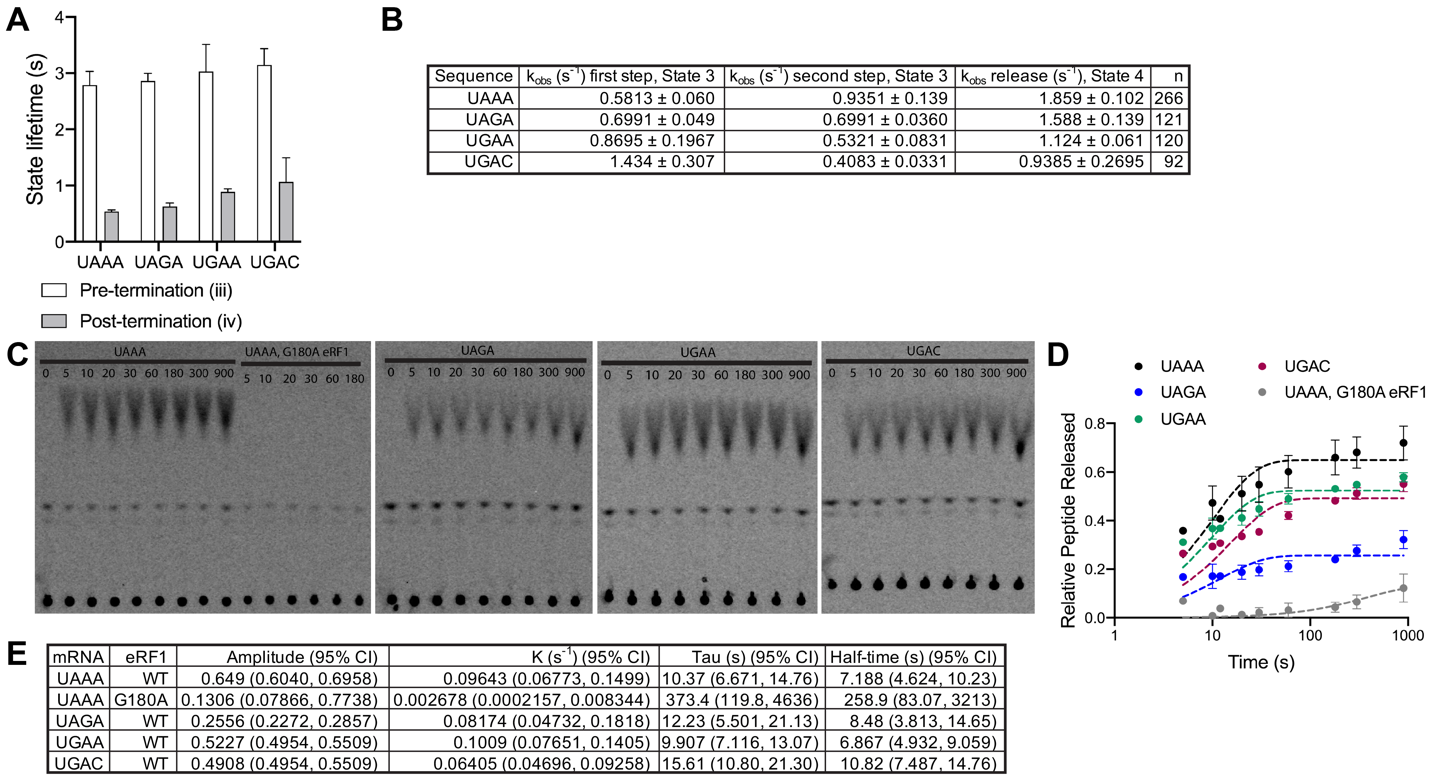

**Fig. S8**: The impact of stop codon sequence on termination kinetics. (**A-B**) Peptidyl-tRNA bond hydrolysis and eRF1 release occur at similar rates with all three stop codons; little impact is observed for the +4 base. (**A**) State lifetimes and (**B**) statistics on fits. (**C-E**) Peptides are released at similar rates with all three stop codons; a subtle effect is observed for the +4 base. (**C**) example TLCs (time in seconds), (**D**) fraction of peptide released with one-phase association fits, and (**E**) fitting statistics.

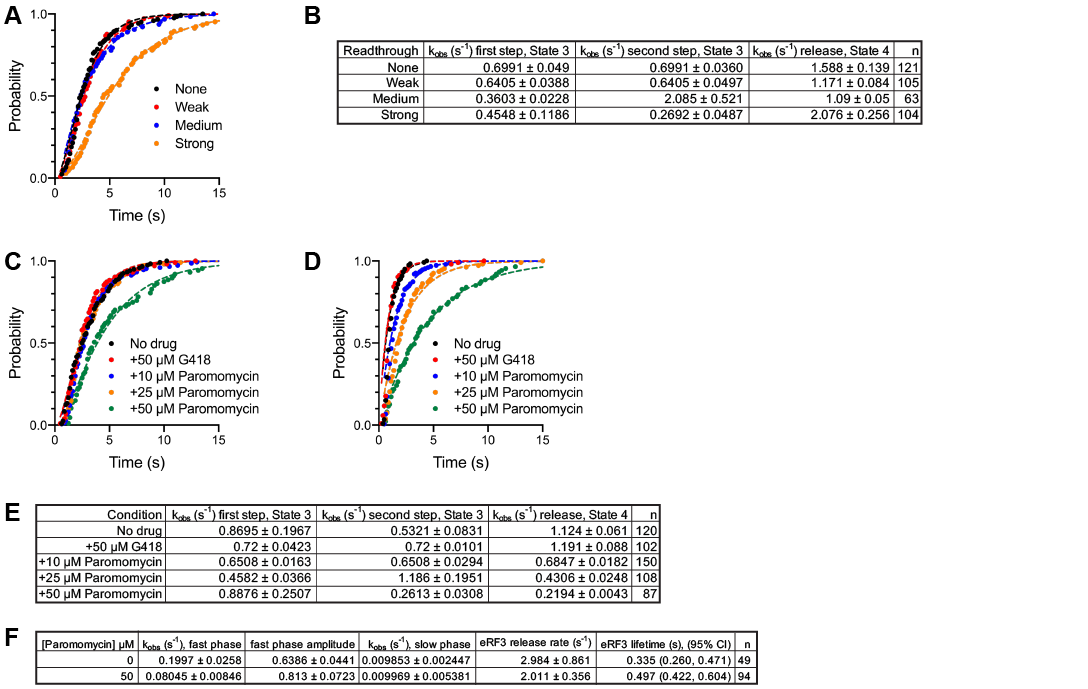

**Fig. S9:** Inhibition of termination by readthrough-promoting 3’ UTR sequences or the aminoglycoside paromomycin. (**A-B**): Readthrough-promoting 3’ UTR sequences inhibit termination. (**A**) Pre-termination state lifetime with fits and (**B**) fitting statistics. (**C-E**): The aminoglycoside paromomycin inhibits termination and eRF1 release; G418 does not affect either. (**C**) Pre-termination state lifetime with fits, (**D**) Post-termination state lifetime with fits, and (**E**) fitting statistics. (**F**) Paromomycin hinders co-binding of eRF1 and eRF3 to M-Stop ribosomes.

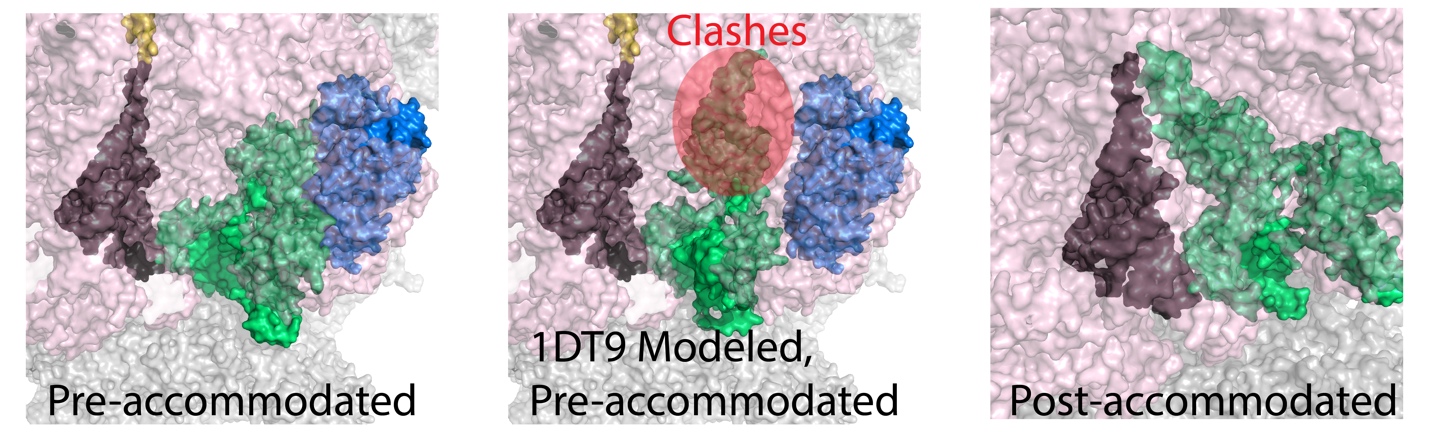

**Fig. S10:** Release factor binding sites on the ribosome. eRF1 shown in green, eRF3 in blue, P-site tRNA in black, and nascent peptide in yellow. *Left*, eRF1 in pre-accommodated state (5LZT, (*14*)). *Middle*, the conformation of eRF1 observed in solution (1DT9, (*26*)) aligned to the N-terminal domain of pre-accommodated eRF1 clearly clashes with the ribosome (highlighted in red). *Right*, eRF1 in the post-accommodated state (PDB ID: 5LZU, (*14*)).

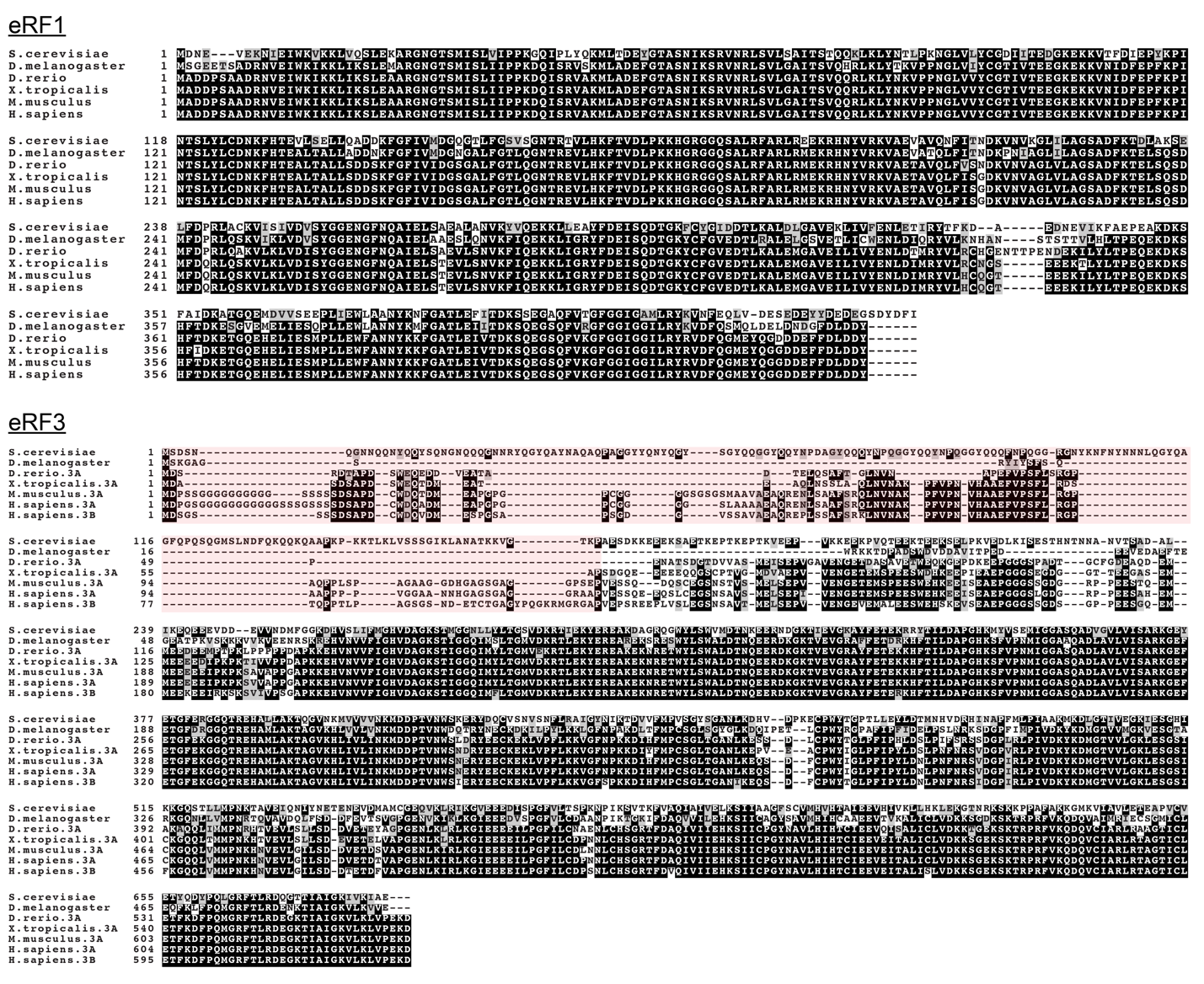

**Fig. S11**: The eukaryotic release factors are highly conserved from yeast to humans. Alignment positions with at least 50% agreement are shaded in black, and the poorly-conserved N-terminal region of eRF3 that was excluded from constructs used in this study (eRF3∆165N) is shaded in light red.

**References (38 – 54)**

38.	J. Yin *et al.*, Genetically encoded short peptide tag for versatile protein labeling by Sfp phosphopantetheinyl transferase. *Proc. Natl. Acad. Sci. USA* **102**, 15815-15820 (2005).

39.	E. Gutierrez *et al.*, eIF5A promotes translation of polyproline motifs. *Mol. Cell* **51**, 35-45 (2013).

40.	C. J. Shoemaker, D. E. Eyler, R. Green, Dom34:Hbs1 promotes subunit dissociation and peptidyl-tRNA drop-off to initiate no-go decay. *Science* **330**, 369-372 (2010).

41.	P. Tesina *et al.*, Molecular mechanism of translational stalling by inhibitory codon combinations and poly(A) tracts. *The EMBO journal* **39**, e103365-e103365 (2020).

42.	J. Wang *et al.*, Structural basis for the transition from translation initiation to elongation by an 80S-eIF5B complex. *Nature Communications* **11**, 5003 (2020).

43.	A. G. Johnson *et al.*, RACK1 on and off the ribosome. *RNA (New York, N.Y.)* **25**, 881-895 (2019).

44.	O. V. Korobov V., *Chemical Kinetics with Mathcad and Maple*. (Springer-Verlag/Wien, 2011).

45.	A. Petrov, R. Grosely, J. Chen, S. E. O'Leary, J. D. Puglisi, Multiple Parallel Pathways of Translation Initiation on the CrPV IRES. *Mol. Cell* **62**, 92-103 (2016).

46.	R. A. Marshall, M. Dorywalska, J. D. Puglisi, Irreversible chemical steps control intersubunit dynamics during translation. *Proc Natl Acad Sci U S A* **105**, 15364-15369 (2008).

47.	C. E. Aitken, J. D. Puglisi, Following the intersubunit conformation of the ribosome during translation in real time. *Nat. Struct. Mol. Biol.* **17**, 793-800 (2010).

48.	M. F. Juette *et al.*, Single-molecule imaging of non-equilibrium molecular ensembles on the millisecond timescale. *Nat. Methods* **13**, 341-344 (2016).

49.	J. E. Bronson, J. Fei, J. M. Hofman, R. L. Gonzalez, Jr., C. H. Wiggins, Learning rates and states from biophysical time series: a Bayesian approach to model selection and single-molecule FRET data. *Biophys. J.* **97**, 3196-3205 (2009).

50.	H. S. Steinert, J. Rinnenthal, H. Schwalbe, Individual basepair stability of DNA and RNA studied by NMR-detected solvent exchange. *Biophys. J.* **102**, 2564-2574 (2012).

51.	J. Rinnenthal, B. Klinkert, F. Narberhaus, H. Schwalbe, Direct observation of the temperature-induced melting process of the Salmonella fourU RNA thermometer at base-pair resolution. *Nucleic Acids Res.* **38**, 3834-3847 (2010).

52.	C. Magis *et al.*, T-Coffee: Tree-based consistency objective function for alignment evaluation. *Methods Mol Biol* **1079**, 117-129 (2014).

53.	P. Saini, D. E. Eyler, R. Green, T. E. Dever, Hypusine-containing protein eIF5A promotes translation elongation. *Nature* **459**, 118-121 (2009).

54.	M. H. Park, H. L. Cooper, J. E. Folk, Identification of hypusine, an unusual amino acid, in a protein from human lymphocytes and of spermidine as its biosynthetic precursor. *Proc. Natl. Acad. Sci. USA* **78**, 2869-2873 (1981).
